# Medieval genomes from eastern Iberia illuminate the role of Morisco mass deportations in dismantling a long-standing genetic bridge with North Africa

**DOI:** 10.1101/2024.10.09.617385

**Authors:** Gonzalo Oteo-García, Marina Silva, M. George B. Foody, Bobby Yau, Alessandro Fichera, Llorenç Alapont, Pierre Justeau, Simão Rodrigues, Rita Monteiro, Francesca Gandini, Marisa Rovira, Albert Ribera i Lacomba, Josep Pascual Beneyto, Valeria Mattiangeli, Daniel G. Bradley, Ceiridwen J. Edwards, Maria Pala, Martin B. Richards

## Abstract

**Background:** The Islamic influence on the Iberian Peninsula left an enduring legacy culturally and linguistically, however the demographic impact is less well understood. This study aims to explore the dynamics of gene flow and population structure in eastern Iberia from the early to late Medieval period through ancient DNA.

**Results:** Our comprehensive genomic analysis uncovered gene flow from various Mediterranean regions into Iberia before the Islamic period, supporting a pre-existing pan-Mediterranean homogenization phenomenon during the Roman Empire. North African ancestry is present but sporadic in late antiquity genomes but becomes consolidated during the Islamic period. We uncovered one of the earliest dated Islamic burials in Spain, which showed high levels of inbreeding. For the first time we also prove the persistence of North African ancestry in a Christian cemetery until the 17th century, in addition to evidence of slave traffick from the Maghreb.

**Conclusions:** This study reveals the complex interaction between political events and cultural shifts that influenced the population of eastern Iberia. It highlights the existence of a slave trade and underscores the lasting impact of historical events, such as the Expulsion of the Moriscos in 1609 CE, on the region’s genetic and cultural landscape through mass population displacement and replacement.

## INTRODUCTION

Andalusi Arabic was still spoken in the Iberian Peninsula in the 17th century CE, with its last remnants surviving in the hinterland of the Kingdom of Valencia (1). This situation of social and linguistic isolation endured by part of the local population was the closing chapter of a story of acculturation that started nine centuries prior. Although this process involved a degree of genetic admixture between newcomers and local Hispano-Romans (2), this admixture is not fully reflected in the present-day population (3).

Several ancient genomes have highlighted patterns of long-distance mobility in antiquity and medieval times across the Mediterranean (4–8). Although some of these admixture events appear to have been transient (5,6), their impact was almost as transformative as previous prehistoric migrations (9,10). Interestingly, however, their extent was more restricted in space, and most noticeable in urban centres (5–8). The field of aDNA has recently started to shift the focus from continental scales to smaller scale, focussed, archaeological questions in European contexts (11–15).

The genetic past of the Iberian Peninsula as a whole has been a subject of extensive research in the last few years, both through modern (3,16,17) and aDNA. However, despite the significant amounts of data produced, the sampling effort has not been homogeneous across periods, due to research interests or sample availability (18), with the majority of studies focusing on prehistory (19–27). The historical periods, although extensively studied by historiography, have only been observed cursorily through the lens of aDNA (2,15,28).

The eastern lands of Spain, which today comprise the Valencian territory, appear to have been a buffer zone between Greek colonies in the northeastern coast and Punic-Carthaginian colonies in the southeast (29). Indigenous settlements did exist in the region, and some even gained temporary notoriety in the early phase of Roman Republican occupation. One such was Arse, later known as Saguntum (modern Sagunto), whose siege by Hannibal Barca in 219 BCE triggered the start of the Second Punic War. Later developments include the foundation of the colony of *Valentia* (modern Valencia) in 138 BCE as one of the earliest outside Italy (30–33).

In the later Visigothic period, this land overlapped with the contested province of *Spania* under Byzantine occupation in the centuries prior to the Islamic conquest (552–624 CE). During this time, Valencia was a semi-autonomous city administered by bishops subordinate to the kings in Toledo (34,35). The territory and the city were surrendered in the Treaty of Tudmir (36–38) by the local Visigothic governor (39,40).

The Islamic conquest (711-756 CE) led to the establishment of Al-Andalus (41) and resulted in centuries of acculturation, co-existence, assimilation, linguistic intertwining and conflict in the Iberian Peninsula.

Al-Andalus was initially shaped by a phase of Arab political dominion under the Umayyads during both Emirate (756-929 CE) and Caliphate (929-1031 CE) periods. Following the collapse of the Umayyad Caliphate, fragmentation ensued. The instability of the petty Islamic kingdoms during the Taifas periods led to two invasions by two North African empires between 1090 and 1237. As a result, Al-Andalus came to be dominated by successive Amazigh (Berber) military powers, the Almoravids (1040-1147 CE) and the Almohads (1121-1269 CE) (41).

The introduction of Islam resulted in a new societal organisation amongst the local Visigothic and Hispano-Roman population (41). In addition to a new Arab minority, many Amazigh tribes came to settle in the Peninsula. This resulted in varied cultures and communities across the Iberian Peninsula: Christians living outside of Al-Andalus in the North, Christians living in Muslim territories of Al-Andalus (*Mozarabs*), and converts to Islam in Al-Andalus (*Muladis*), who came to represent the majority of the population.

By the end of the 13th century CE, the Christian kingdoms controlled the majority of Iberian territory. However, the *Muladis* still accounted for a large fraction of the population, and remained so instrumental to the economy in certain regions that the new Christian elites allowed them to maintain their cultural and religious traditions (42), thus becoming the *Mudéjar*, *i.e.* Muslims living in Christian territories (43,44). However, by the 16th century CE, religious tensions against converts and *Mudéjars* had increased. As a result, in the first half of this century, the *Mudéjars* became targets of forced mass conversions, and the converts to Christianity became known as *Moriscos* (45).

In practice, the *Moriscos* kept their language and customs, and many continued practising their religion in secret (45). This was particularly true for *Moriscos* in the Kingdom of Valencia (1,42,46,47). However, this dualism came to an end in 1609 CE, when Phillip III decreed the expulsion of all *Moriscos*. Estimates suggest that, in the Kingdom of Valencia, more than one third of the total population was expelled to North Africa (48). The total numbers are estimated to have amounted to more than 300,000 people (49,50).

Altogether, these historical episodes highlight the possibility for the arrival, admixture and dwindling of varied sources of ancestry in different historical periods (2,4). Here we investigate the extent to which North African-related ancestry fluctuated in eastern Iberia during historical times, with particular focus on the genetic impact of the establishment of Al-Andalus and later conquest of the region by Christian kingdoms (51). For this end, we generated shotgun genome data from 12 individuals from various locations across the eastern Mediterranean coast of Iberia, including three pre-Islamic burials (from Roman and late Visigothic periods), five Islamic inhumations, and four individuals from a Christian cemetery (both medieval and post-medieval), covering a period of over one thousand years.

## RESULTS

### Time transect overview and uniparental diversity

We generated new historical genomes from 12 individuals (Table 1; Figure 1A,B,C) from three sites in the Valencian region (Figure 1D). The time transect (Figure 1E) spans late antiquity (*n* = 3), medieval Islamic (*n* = 5), late medieval Christian (*n* = 2), and post-medieval Christian (*n* = 2) stages. This set of individuals contributes the first shotgun-generated historical transect for Spain (52).

**Figure 1:**
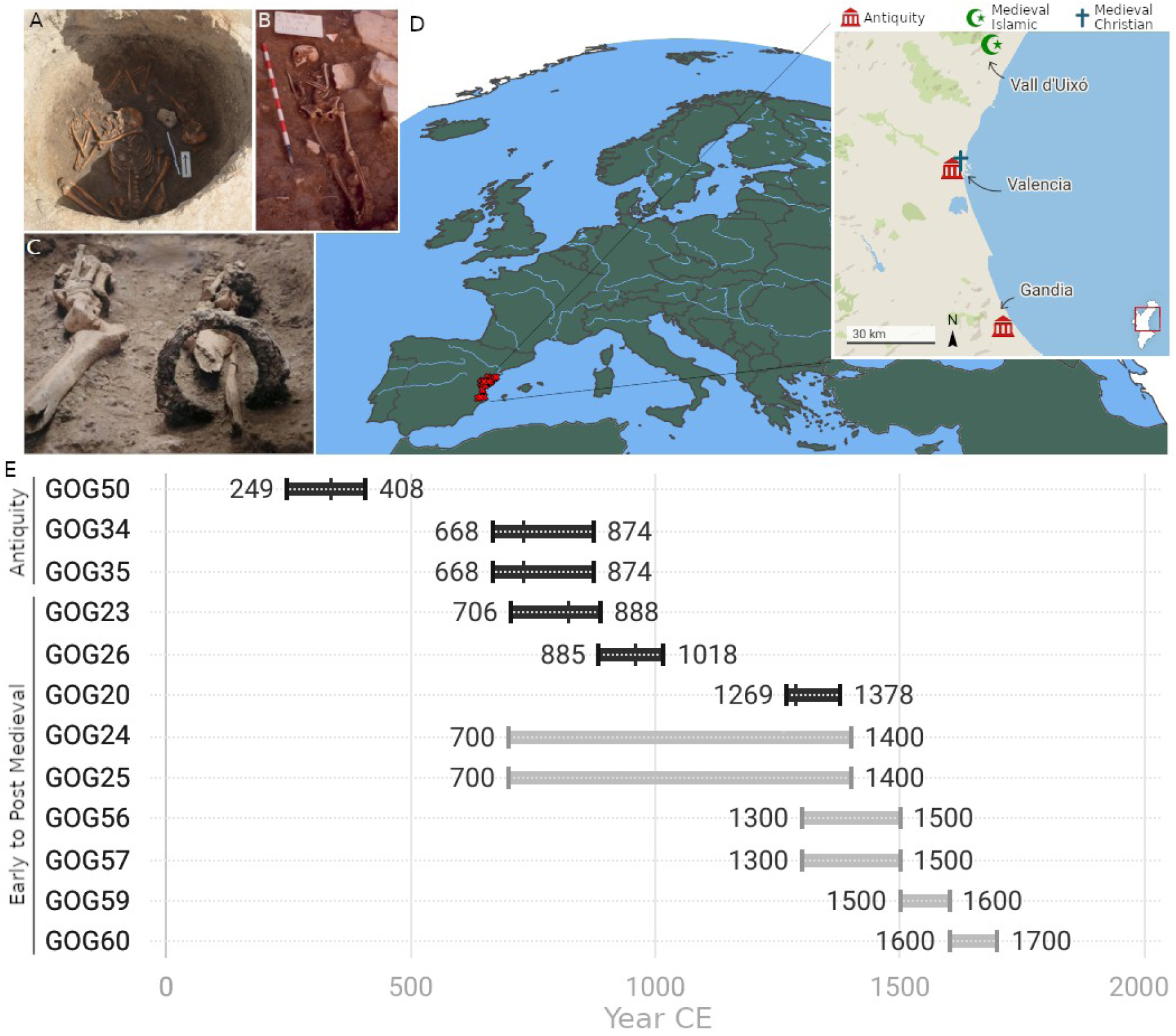
A) Two kindred individuals from late antiquity contemporary to the Islamic invasion found in Gandía. B) Early Islamic burial (GOG26) from the 8th century CE in Vall d’Uixó, sequenced to 2.3X coverage. C) Remains of sample GOG59 from Valencia, consisting solely of the lower extremities and an iron shackle on the right leg. D) Map with the location of all samples screened (red crosses) in this Mediterranean region. The minimap details antiquity (Roman and post-Roman), Islamic and Christian (Late and post-Medieval) sites where the newly sequenced samples in this study originate. E) Timeline with the dates for each sample studied. Samples with bars in black indicate calibrated radiocarbon dates, internal ticks indicate median calibrated age, and the range indicates 95% confidence intervals, OxCal version 4.4.4 and IntCal20 (Bronk Ramsey 2009; Reimer et al., 2020). Light grey bars indicate relative dates based on stratigraphy, material culture or context from excavation.

**Table 1:** Overview of individuals sequenced in this study.

| Lab ID | Location | Period, culture | Radiocarbon dates | Genetic sex | mtDNA haplogroup | <sup>1</sup>ChrY haplogroup | <sup>2</sup>Genome coverage |
| --- | --- | --- | --- | --- | --- | --- | --- |
| GOG5 | Valencia, | Antiquity, | 249-408 | Femal | D4e1 | - | 1.88x |
| 0 | San Lorenzo | Roman | (cal.) CE | e |  |  |  |
| GOG3<br>4 | Gandía,<br>Sanxo<br>Llop | Antiquity,<br>Post-Roman | 668-874<br>(cal.) CE | Male | HV+1631<br>1 | R1b-L11 | 0.26x |
| GOG3<br>5 | Gandía,<br>Sanxo<br>Llop | Antiquity,<br>Post-Roman | 668-874<br>CE* | Femal<br>e | H2a1e1a | - | 0.36x |
| GOG2<br>4 | Vall<br>d'Uixó,<br>Benigafull | Medieval,<br>Islamic | - | Male | U4a1d | J2a-M260 | 0.51x |
| GOG2<br>5 | Vall<br>d'Uixó,<br>Ceneja | Medieval,<br>Islamic | - | Femal<br>e | L3d1 | - | 0.15x |
| GOG2<br>3 | Vall<br>d'Uixó,<br>Benigafull | Medieval,<br>Islamic | 706-888<br>(cal.) CE | Male | HV | E1b-PF2477 | 0.63x |
| GOG2<br>6 | Vall<br>d'Uixó,<br>Ceneja | Medieval,<br>Islamic | 885-1018<br>(cal.) CE | Femal<br>e | H1 | - | 2.34x |
| GOG2<br>0 | Vall<br>d'Uixó,<br>Benizahat | Medieval,<br>Islamic | 1269-1378<br>(cal.) CE | Femal<br>e | J1c1b | - | 0.47x |
| GOG5 | Valencia, | Late | - | Femal | H+7720 | - | 0.21x |
| 6 | San Lorenzo | Medieval, Christian |  | e |  |  |  |
| GOG57 | Valencia, San Lorenzo | Late Medieval, Christian | - | Female | R0a4 | - | 0.17x |
| GOG59 | Valencia, San Lorenzo | Post-Medieval, Christian | - | Male | H5+152 | E1b-5081 | 0.26x |
| GOG60 | Valencia, San Lorenzo | Post-Medieval, Christian | - | Male | K1a+195 | R1b-L483 | 0.23x |

<sup>1</sup>Y chromosome classification as per ISOGG 2019 – most derived SNP.
<sup>2</sup>Coverages reported in this table calculated after mapping quality (MQ >20) and read length (bp >34) filters applied.
\*By association with GOG34, same layer but not directly dated.

We selected these twelve individuals on the basis of DNA preservation from the total of 35 samples screened, which yielded a range of endogenous DNA proportions from 0.01 to 41.58%, with the average endogenous DNA content being 7% (calculated as the proportion of mapped reads longer than 34 bp with mapping quality >30) (Figure S1). The sequencing of the DNA recovered from these 12 samples, which included petrous bones, molars and a metatarsus, yielded average genome depth coverages between 0.21× and 2.34× (Table 1, Figure S1).

We also report new radiocarbon dates (Table 1, Figure S11) for three burials from antiquity and three medieval Islamic burials. To the best of our knowledge, the new dates uncovered one of the earliest Islamic burials with genomic data ever found in the Iberian Peninsula (GOG23, Figure 1B). For the remainder of samples without radiocarbon dating we used relative dating, based on archaeological context, material culture and other information from excavation reports. The twelve individuals form the scaffold form a transect that starts in the third century CE and stretches to the seventeenth century CE (Figure 1E).

The remains of the Roman burial (GOG50) from the city of Valencia had good endogenous DNA preservation (35%) in the context of this study. We identified the genetic sex of this individual as female (Tables S1, S15), carrying the mtDNA lineage D4e1. Notably, haplogroup D4e1 belongs to an East Asian clade that is rare in Europe throughout time (53,54). The radiocarbon dating of the locates the sample between the years 249–408 cal. CE (median 338 CE), is in line with its archaeological context. These dates indicate the woman lived through a period of intense change in the Roman world marked by the Christian persecutions and the Edict of Milan (313 CE). However, although poorly preserved (52), the burial appears pagan in nature as it was accompanied by the remains of a swine head. Such offerings are also found in much earlier burials from the Republican period as part of the *porca praesentanea* ritual (55).

The remains of an adult (GOG34) and an infant (GOG35) from the late Visigothic period in Spain (668–874 cal. CE) (Figure 1A, Table 1) were found together in a round pit excavated in a site near Gandía (Figure 1A,D). Genetic analysis identified the adult (GOG34) as male and the infant (GOG35) as female (Table S15). They carried mtDNA haplogroups HV+16311 and H2a1e1a, respectively. The adult carried a R1b1a1b1a1a-L11 (likely unresolved) Y-chromosome (chrY) haplogroup. READ (Figure S3) detected a first-degree kinship between these two individuals. Combined with their ages at death, simultaneous burial, sex identification and non-matching mtDNA haplogroups, we inferred the kinship status to be father–daughter (56).

From the medieval and later periods we sampled various sites from Vall d’Uixo and the city of Valencia. Samples from Vall d’Uixó belonged to various Islamic cemeteries (*Maqbaras*) of rural settlements in the higher (Alquerías Ceneja and Benifagull) and lower (Alquería Benizahat) areas of the modern town (57). As they expanded, these settlements eventually fused together (57). We obtained older radiocarbon dates (706–888 and 885–1028 cal. CE, respectively) for individuals from the higher settlements (GOG23 and GOG26; Figures 1E, S11, Table 1), whereas the individual from the lower settlement was dated to a younger age (1269–1378 cal. CE). Some of these radiocarbon dates are much older than anticipated but confirmed a suspicion among local archaeologists that the high ground settlement had been the original.

Among these samples, we found west Eurasian mtDNA haplogroups U4a1d, HV, H1 and J1c1b, as well as the more typically African haplogroup L3d1. As for the chrY lineages we found J2a1a1a2b2a1a-M92 and E1b1b1b1a1-M183 in the two males (Table 1). J2a1a1a2b2a1a-M92 likely has a Levantine origin (58,59) but was present in Italy by Roman times (6). E1b1b1b1a1-M183 is overwhelmingly the most common extant male lineage in North Africa, where it expanded within the last 3000 years (60) (Tables 1, S16).

The burials from a medieval Christian cemetery that remained in use until the mid-19th century CE in Valencia, were dated based on archaeological context and historical record. Two of these (GOG56 and GOG57) were assigned as late medieval graves from the 14th-15th centuries CE. We identified both these individuals as genetically female, with mtDNA haplogroups H+7720 and R0a4. The others (GOG59 and GOG60) were post-medieval graves of two males dating to the 16th and 17th century CE respectively. From the 16th century individual (GOG59), as seen in Figure 1C, only the lower body was recovered with an iron shackle around the bones of his right leg still visible, suggesting enslavement. This individual carried a West Eurasian H5 mtDNA lineage and a North African E1b1b1b1a1-M183 chrY lineage. GOG60 carried a West Eurasian K1a mtDNA and an unresolved West Eurasian R1b1a1b-M269 chrY lineage (Table 1).

### Pan-Mediterranean genomic homogenization trends in the eastern coast of Roman Hispania

All three samples from antiquity appear as significant outliers with respect to the modern Spanish population in the PCA (Figure 2A).

**Figure 2:**
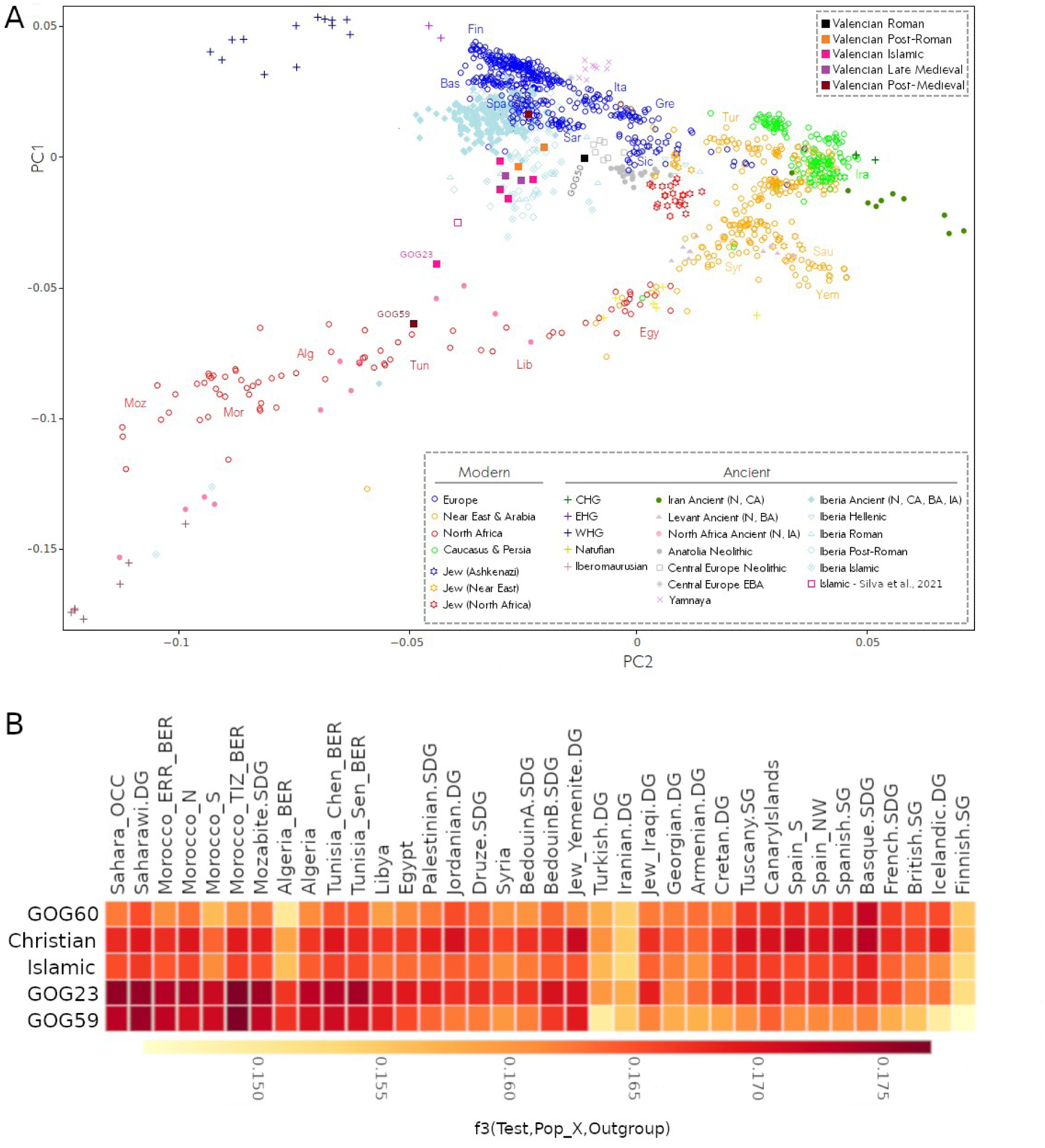
A) PCA plot built with modern populations from around the Mediterranean world on top of which key ancient populations and newly sequenced genomes from this study were projected. B) Outgroup-f3 test measuring shared drift of the groups defined by the ancient Valencian samples (Test) in this study with North African, Levantine and European modern populations (Pop_X). The Islamic grouping includes GOG20, GOG24, GOG25 and GOG26. The Christian grouping encompasses GOG56 and GOG57.

The Roman female individual (GOG50) trends towards modern central and eastern Mediterranean populations, sitting midway between present-day Spanish and Sicilians (Figure 2A). Her heterozygosity levels are higher however, suggesting multiple ancestry contributions (Table S12, Figures S7, S8, S10). In the PCA, this individual shares a hybrid space defined by some other Roman Iberian and Punic individuals from Sardinia (8). We conclude that her affinities were derived from a complex admixture process of two genetic sources that do not exist today as consolidated populations. One parental source was Iberian, but with higher North African ancestry than present-day Spaniards (2,3), and the other was Italian-Sardinian, with eastern Mediterranean ancestry, as seen in the Roman Imperial period (6). However, it is difficult to pinpoint the exact sources, due to the extremely high heterogeneity observed among individuals in the Mediterranean during late Roman times (6–8). Proximal *qpAdm* modelling also highlights the contribution of an eastern Mediterranean source to her ancestry (Table S13).

Furthermore, the mitochondrial haplogroup D4e1 (61) carried by GOG50 is extremely rare amongst ancient Europeans and Near Easterners. There are only a couple of instances in present-day Europeans. Modern data show that D4 and D4e lineages (except D4e1) are all found in East Asia. Other D4e1 sister haplotypes are also widely dispersed in Central, East and Southeast Asia and, to a minor degree, in South Asia (54).

In particular, haplogroup D4e1 has been identified in a Xiongnu individual (TAK008) from the Altai, dating to 51–155 BCE (62), whilst it is not seen amongst 60 Xiongnu individuals from further east (63). The Xiongnu formed a diverse steppe empire, which might be a potential source for the lineage in Europe through third-century CE dispersals of Huns, Alans and Sarmatians (64). It has also been identified in a burial of similar age (203–319 cal. CE) from the Roman era in Turkey (59). Our finding of the lineage in Iberia, so far west in the Mediterranean, seems strikingly early. However, as shown in (11), migrants from Eastern Europe were already passing through Hispania before the start of the so-called Germanic Invasions.

To trace recent East Asian ancestry in the genome of this individual we made use of *f*-statistics and LAI. None of the *f*-statistics detected extra genetic affinity to Asian populations (Table S12). Similarly, LAI evaluation with RFmix did not reveal outstanding stretches of East Asian-related ancestry but found some minor South Asian-like haplotypes (Figure S9). This result mirrors the finding that the Xiongnu D4e1 individual (TAK008) already carried around 90% west Eurasian ancestry (62). The two kindred individuals from late antiquity (GOG34 and GOG35, father and daughter, respectively) occupy a space with some other post-Roman and medieval Islamic samples from Iberia, also shifted towards the western end of the North African cline of the PCA. ADMIXTURE and *qpAdm* results indicate that the father (GOG34) carried North African-related ancestry (tested using *Iberomaurusian* and *Morocco_EN* as proxies, respectively) (Figure 3A, Tables S13, S14). However, in ADMIXTURE the daughter (GOG35) displays approximately half the amount of such ancestry than the father. This likely indicates that the mother contributed none and suggests different genomic backgrounds of the two parents. In fact, *qpAdm* models including sources with such ancestry are rejected for the daughter (GOG35) but not by LAI analysis. The case of GOG34 is interesting since his genome-wide pattern aligns with the core of Islamic genomes and his median C14 dating (734 cal. CE) makes him contemporary with the start of Islamic conquest – yet the archaeological context appears pre-Islamic.

**Figure 3:**
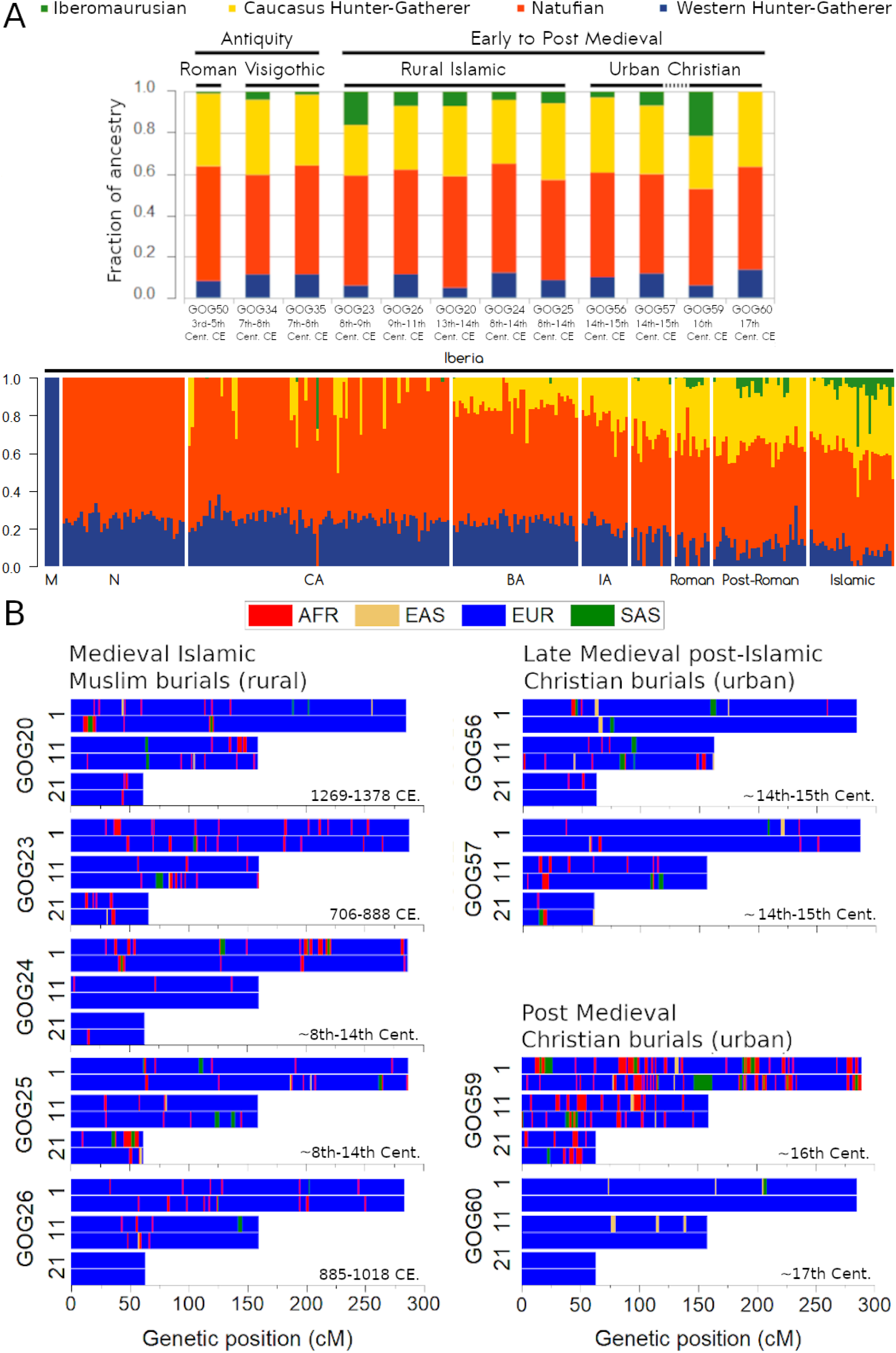
A) Supervised ADMIXTURE (K=4) defining a distal model, using as proxy sources for ancestry determination differentiated hunter-gatherer groups from Europe (WHG), Caucasus (CHG), the Levant (Natufians) and North Africa (Iberomaurusians). B) RFmix chromosome ancestry segments coloured according to four possible sources from the 1000 Genomes Project: African (AFR) as proxy for North African contribution, European (EUR) representing the Iberian contribution, East Asian (EAS) and South Asian (SAS) as proxies for broad Asian ancestry.

Supervised and unsupervised ADMIXTURE, *qpAdm* models (Figures 3A, S6, Tables S13, S14) and LAI results (Figures 3B, S9) confirm that all pre-Islamic individuals carried North African ancestry at lower levels than the later medieval Islamic individuals. In pre-Islamic times the influx of North African-related ancestry was more sporadic, since it is not found consistently across all published individuals. It is only with the onset of the Islamic period that it becomes more sustained and consolidated at a higher percentages (Figure 3A).

### Survival of widespread North African-related ancestry until the 17th century in the Valencian kingdom

The group of medieval Islamic individuals (GOG20, GOG23, GOG24, GOG25, GOG26) appears homogeneous in the PCA (Figure 2A) with the exception of GOG23. They form a core cluster together with pre-Islamic GOG34 and two individuals (GOG56 and GOG57) from the late medieval post-Islamic period (Figure 2A), in addition to some other published Islamic individuals from elsewhere in Spain (2). We identify this cluster space as a consolidated ‘*Berberized’* population that formed largely during the Al-Andalus era (SI Figures S5, S6). In our region of study, North African-related ancestry is present in these individuals at ∼14–18% estimated with *qpAdm* (with SEs <3% and *p*-values ranging from 0.43 to 0.89). These are slightly higher levels than in the earlier individuals, as seen in the supervised ADMIXTURE (Figure 3A). GOG23 is the oldest Islamic burial dated in our dataset (706–888 cal. CE) and constitutes, to our knowledge, one of the earliest examples of an Islamic burial from eastern Iberia to be studied genetically. This individual is drawn even closer in the direction of North African populations in the PCA (Figure 2A), and had the highest amount of North African-related ancestry, as shown by the ADMIXTURE (Figure 3A), *qpAdm* (∼31±2.5% (±1 SE); *p*=0.13) and LAI (Figures 3A, 3B). Using temporally proximal sources, we could model GOG23’s ancestry as deriving ∼50% from a source genetically resembling *CanaryIslands_Guanche.SG* and ∼50% from a source similar to *Portugal_LateRoman.SG* (*p*-value=0.09; SE=4.9%) (Tables S13, S14).

The two late medieval samples (GOG56 and GOG57) from the 14th–15th century CE, were recovered from a Christian cemetery belonging to the parish of San Lorenzo in the city of Valencia. However, these two individuals still group within the medieval ‘*Berberized’* PCA cluster (Figure 2A) two centuries after the Christian conquest of the city. The levels of North African-related ancestry are still comparable to those observed in the Islamic period (Figure 3, Tables S13, S14).

Post-medieval individual GOG59, who was found with an iron shackle around his right ankle clusters closely with modern Moroccans and Algerians in the PCA (Figure 2A), indicating a North African origin. Most present-day Berber individuals in the North African PCA fall along a cline defined by the amount of sub-Saharan ancestry. GOG59 sits at the low sub-Saharan ancestry side of this cline (Figures S4, S5). Outgroup-*f3* analysis shows that the highest shared drift is with present-day Moroccan Berbers (Figure 2B). We can model ∼40 to 50% (SEs ∼3%) of his ancestry as deriving from a North African-related source, depending on the model tested (Tables S13, S14). This result is further supported by a second PCA computed using an alternative Affymetrix North African dataset where GOG59 appears closest to Berber individuals from Morocco and Algeria (Figures S4, S5).

In contrast, the other post-medieval individual from the same cemetery, GOG60 (Figure 2B), was the only one in our dataset that fully clustered in the PCA within the modern Spanish population (Figure 2A). No traces of North-African ancestry could be identified in his ADMIXTURE profile or *qpAdm* analyses (Figure 3A, Tables S13, S14). The LAI profile (Figure 3B) also indicated no discernible trace of sub-Saharan African haplotypes in the tested chromosomes. Furthermore, neither uniparental marker indicates affiliations to the Maghreb, but instead rather typical European lineages (Table 1). This profile is in sharp contrast with GOG59, who appears to be of full North African origin. It also contrasts with the immediately preceding individuals from the late medieval period (Figures 2, 3) – whether Muslim or Christian – and the three individuals from antiquity (Figure S9).

### Potential genomic impact of social isolation of Mudéjar and Morisco communities

Based on the information gathered in the analyses above, we grouped some samples into populations (GOG20, GOG24, GOG25, GOG26 as “medieval Islamic” and GOG56 and GOG57 as “Late Medieval Christian”), whilst treating the PCA outliers individually (GOG59, GOG60 and GOG23). Furthermore, by combining the dates (Figures 1C, S11), genetic ancestry, and burial context, we argue that we can identify some individuals as belonging to different socio-historical groups: GOG23 and GOG26 fall into the *Muladi* category (Christian converts to Islam), GOG20 into the *Mudéjar* (Muslims in Christian territories) and GOG56 and GOG57 could be identified as *Morisco* (Muslims forcibly converted to Christianity) or descendants thereof.

To measure the evolution of genetic affinities of our archaeological samples with modern populations from Iberia and North Africa, we ran outgroup-*f3* tests by merging populations from the Human Origins and (65) datasets (Figure 2B). The results obtained follow what we observed in the PCA, ADMIXTURE, *qpAdm* and LAI. The measurements of shared drift indicate a trend of increased affinity towards modern Iberian groups as the samples become more recent with the striking exception of the slave, GOG59 (Figure 2B). Both GOG23, who was an early Islamic individual with some Iberian ancestry, and GOG59 who seems fully North African in ancestry, displayed the highest outgroup-*f3* values with present-day Maghrebi populations, especially Moroccan Berbers from Tiznit (Morocco_TIZ_BER) (Figure 2B). The Islamic core group of four individuals (GOG20, GOG24, GOG25, GOG26) do not show particularly high affinities with either North African/Berber groups or Iberians/Europeans, highlighting more balanced contributions of both sources to their ancestry (Figure 2B) in relation to the other samples analysed. The late medieval Christian grouping was also similar, although with signs of increased affinity with present-day Basques (Figure 2B). Finally, GOG60, the post-medieval individual from the 17th century, displayed the opposite pattern to GOG59 and GOG60, with highest *f3* values shared with Iberian groups (Figure 2B).

Due to coverage and sample size constraints, in order to study ROH patterns in depth we had to combine two approaches: hapROH on the whole-genome and oROH/pROH (see Methods) from the three imputed chromosomes. The use of hapROH allowed us to explore all chromosomes but with loss of individuals. With imputation we evaluated all individuals but in a subset of chromosomes. We validated the concordance of both approaches through the identification of the same ROH fragment in chromosome 21 of individual GOG20 (Figures S7, S12).

In the context of modern worldwide populations, none of the ancient individuals analysed seemed to suffer from extreme levels of consanguinity based on the number of pROH fragments found in the imputed chromosomes (Figure 4A). Three of them (GOG20, GOG34, GOG60) appear on the fringes of normal European variability (Figure 4A). However, hapROH identified the early Islamic burial (GOG23) as highly inbred (Figure 4C). We missed this pattern with oROH since the chromosomes picked do not display noticeably long homozygous segments (Figure S12).

**Figure 4:**
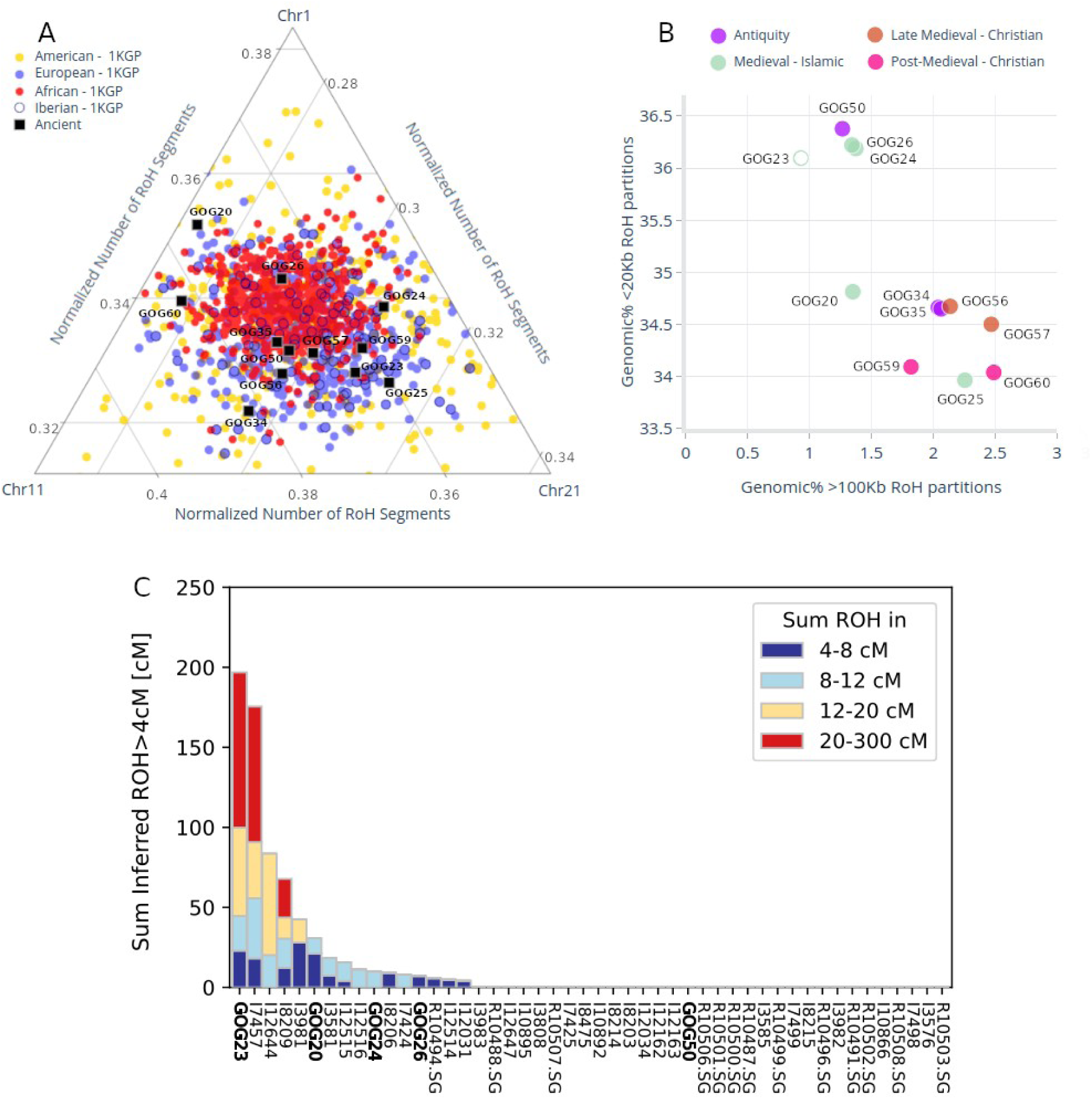
A) Number of runs of homozygosity (ROH) identified by PLINK and normalised by the length of each chromosome. B) Genomic fraction across the three imputed chromosomes covered by short (<20Kb) and long (>100Kb) segments of organic ROH. GOG23 not filled to denote unreliability. C) hapROH results for a subset of the new samples reported in this work along with a set of public samples from contemporary periods.

Bearing this caveat in mind, we attempted to explore the patterns in finer detail. A deeper look within the set of ancient genomes analysed here suggests increased inbreeding over time in the medieval population of Valencia (Figure 4B). Using the oROH from the imputed chromosomes as reference (Figure 4B), the Roman individual (GOG50) together with the two Islamic genomes (GOG24 and GOG26) appear to have the highest percentage of the genome in short organic ROH (20Kb >35%) and the lowest of long organic ROH (100Kb <1.5%) (Figure 4B). This group also included GOG23, but we excluded it from this inference, since hapROH revealed high inbreeding in the rest of chromosomes (Figure 4C).

The remaining group of medieval individuals are more recent burials, except for GOG25, whose exact age is unknown. In this second group, samples exhibited lower percentages of short oROH (<35%) and a higher fraction of long oROH (>1.5%) (Figure 4B). The only exception was GOG20, the Islamic sample with the most recent radiocarbon date (1285 cal. CE. median age) whose fraction of long ROH was also below 1.5%. The two kindred individuals from late antiquity (GOG34 and GOG35) are more akin to each other and group with the later Islamic and medieval samples, rather than with the late Roman female individual (GOG50).

## 4 DISCUSSION

The results presented here buttress the idea that gene flow from North Africa into Iberia was ongoing in the centuries preceding the Islamic period. However, we also show that this gene flow was not restricted to southern Iberia (2), but it also impacted eastern Iberia since at least late Roman times. This inflow is evidenced by the finding of North African ancestry in all pre-Islamic genomes, albeit at low levels. Our data also suggest that, during the centuries of Roman Imperial rule, there was a significant dynamic of pan-Mediterranean homogenization contributing sporadic Asian-related ancestry, as exemplified by individual GOG50. With lesser intensity, this mirrors the dynamic observed in Rome itself (6), a phenomenon most likely driven by mobile peoples from Italy, Greece, Asia Minor and the Eastern Provinces as slaves or otherwise. However, more genomes are necessary for a full comparison with other localities. This transient phenomenon hints at a much more complex picture if we wish to understand and quantify accurately the genetic contribution of North African migrants in the later Islamic era. We also highlight the apparent lack of ancestry contribution from native peoples from the Arabian Peninsula during the Islamic period in the genomes studied in this work.

Note that a shortcoming of studies focusing on the Islamic period is the lack of contemporary North African and Arabic ancient genomes. A gap that should be filled in the future.

The profile of the genome recovered from the burial of the female (GOG50) from the Roman colony of Valencia provides interesting insight into the heterogeneity of the Roman population of the city. The local Iberian population in coastal regions may have been no stranger to North African influences, which, coupled with shared forms of Romance languages, could have been a factor aiding the later swift Islamic conquest. This is supported by the fact that the two post-Roman individuals also carried North African-related ancestry. The Islamic conquest of the Iberian Peninsula, beginning in 711 CE, further intensified this phenomenon by intertwining the Maghreb and parts of Iberia intimately together (41), also seen at the genetic level (15).

The genomes from the Islamic period therefore point to an intensification and consolidation of the genetic *ńBerberizatioń* in eastern Iberia during the Middle Ages, rather than a completely new phenomenon. The genetic structure of medieval Iberia became a patchwork resulting from the shifting religious and cultural geographies.

We argue that the Islamic burials who carried a higher North African contribution in their genomes were likely representatives of the *Muladi* population: people of Iberian descent who adopted Islamic culture and religion and inter-married (the original meaning of *muladi* in Arabic was a person of mixed Arab and non-Arab ancestry (66–68). The oldest Islamic individual presented here, GOG23, lived within the first generations following the conquest, and had North African and an Iberian-related ancestries, suggesting that acculturation was relatively swift and accompanied by admixture between locals and newcomers. However, his high level of inbreeding might indicate a limited pool of early Muslims. Alternatively, just a mere atypical family tree. Although highly-inbred individuals occur later, in the 12^th^–13th century CE (2), his high levels of inbreeding seem to be somewhat exceptional (Figure 4C). Refined dating and more ancient genomes at higher coverages, as well as contemporary genomes, would be necessary to identify any possible trends in early Islamic burials. On the other hand, the two late medieval Christian burials from Valencia, who carried similar fractions of North African-related ancestry, may be representatives of the *Moriscos*, the people who converted to Christianity at the beginning of the 16th century CE. Mass conversions would explain the obvious genetic continuity, despite the alleged Muslim population displacements and socio-political changes experienced following the Christian conquest.

The apparent trend of growing homozygosity through the medieval period – genomic profiles with similar ancestry composition have more longer organic ROH and fewer short ones in early medieval individuals than in later medieval samples – could be the product of *“habitat fragmentation”* by means of loss of connectivity, social isolation and reduction of *Mudéjar* and *Morisco* communities in the new Christian society. We caution that this observation can only be speculative given the small sample size and should be confirmed with new radiocarbon dates and genomes. Regarding the post-medieval individual GOG60, the reasons for his levels of homozygosity must be found elsewhere. He might originate from another isolated social group of Christians in Islamic society like the *Mozarabs*. Our results also shed light on urban dynamics in Valencia. Despite property seizure by the new Christian elites (69), historically described expulsion of native Muslim citizens, and re-population with Aragonese and Catalan people, the genetic make-up inside the cemetery of San Lorenzo (Valencia) did not change compared to the preceding rural Islamic population from Vall d’Uixo and with other urban Islamic individuals from Valencia (2,15).

It is striking that our sole 17th century genome, GOG60, is the only individual lacking North African-related ancestry. Although it is only a single sample, this result is consistent with the low fraction of North African ancestry found in the modern population of the Valencian territory (3,16,70). We provisionally interpret this pattern as a reflection of a historical event that occurred in 1609 CE: the Expulsion of the Moriscos from the Kingdom of Valencia. This expulsion, and repopulation with peoples from northern territories, effectively erased the local North African component. Christians, especially those from more northern regions, likely carried less North African-related ancestry than the Moriscos that they replaced (3,15), although this remains to be tested. It must be noted that individuals without North African ancestry, such as GOG60, may have also lived in Valencia before the 17th century.

On the other hand, the presence of an enslaved male individual (GOG59) of Berber origin, buried in a Christian cemetery, highlights the lower status of inhabitants with links to the Barbary Coast following the *Reconquista* and before the final expulsion. The elucidation of the genetic origin of GOG59 offers an insight into the provenance of some of the slaves during the Valencian Golden Age (14^th^–16th centuries CE), and substantiates historical accounts of Christian raids to capture slaves in North Africa (71), whilst historical records indicate that the provenance of slaves was not limited to North Africa (72–74).

One final point, highlighted by the survival of North African-related ancestry in substantial proportions until the 17th century, is the widespread presence of such ancestry in present-day South Americans (75). Christian converts were forbidden to migrate to the Americas, although clandestine journeys probably occurred. However, the Maghrebi ancestry signature seen today in South America is too high to be satisfactorily explained by sporadic movement. The high estimates of North African ancestry in South America suggests that colonial migration involved people carrying higher levels of this ancestry than the average in present-day Spain (3,75). Furthermore, the time estimates since the Maghrebi admixture in South America are consistent with the Iberian admixture episode (75). This strongly suggests that most of this ancestry was introduced by the initial colonial immigrants. The two late medieval individuals from Valencia further support this observation: a population with increased Maghrebi ancestry existed at the time in Spain, likely not only in Valencia. Given that cities in the south, such as Sevilla and Cádiz, were the main ports for the colonial voyages to America, we hypothesise that North African-related ancestry also survived in southern regions after the end of the Islamic period and became the source the Maghrebi ancestry introduced in South America.

## 5 CONCLUSION

Our aDNA results suggest a major impact on the genetic structure of the eastern Iberian population as a direct result of an episode of ethnic cleansing and cultural genocide in the early 17th century. Under Philip III, an estimated 300,000 Moriscos, or a third of the population, was expelled to North Africa from across Spain – a much larger figure than even the Jewish expulsions after 1492. These were people, largely of mixed Iberian and North African ancestry, who had been settled in Iberia for hundreds of years, and who had been forcibly converted from Islam to Catholicism a century earlier following the *Reconquista*. Despite a hugely civilising legacy on the culture of Europe, particularly of Iberian Peninsula, that endures to the present day, the *Moors* of Spain were largely dispersed across the Mediterranean between 1609 and 1614 CE. As a result, a genetic bridge between Europe and Africa that had been in place for centuries, and whose legacy we can detect in the genomes of those living in the Valencian region up until the 17th century, was thoroughly dismantled.

## MATERIALS AND METHODS

### Sampling and sequencing

We handled and processed archaeological samples for aDNA extraction, in a dedicated aDNA facility at the University of Huddersfield. A we processed a total of 35 samples, comprising a range of different skeletal elements: petrous bones, molars, phalanxes and a metatarsus. We followed established protocols specific for DNA extraction from ancient remains (76–78). Library preparation followed the (79) protocol with modifications (80,81).

We screened 35 double-stranded (ds) DNA libraries (without post-mortem damage/USER™ treatment), one per sample, on the Illumina MiSeq platform at Trinity College Dublin to evaluate endogenous DNA content (Figure S1). We confirmed aDNA post-mortem damage patterns in the screening non-USER™-treated libraries with mapDamage (v.2.0.7). We then selected the most promising samples and generated USER™-treated dsDNA libraries for further paired-end sequencing on the Illumina HiSeq 4000 platform at Macrogen (Seoul, South Korea) (Figure S2).

### Data processing

We evaluated paired-end read quality with FastQC (v.0.11.558) (82), removed adapters and merged paired-end sequencing reads using leeHom (Renaud et al., 2014) with the flag –ancientdna activated. We mapped reads against the human genome reference hg19 with rCRS as mitochondrial reference using BWA aln and samse commands (v.0.7.5a-r40560) with specifications (-l -n 0.01, -o 2) (83). We performed quality control of the resulting BAM files with QualiMap (v.2.262) (84). We removed duplicated reads using the "rmdup" command in Samtools (v. 1.16) (85). To avoid SNP miscalls due to post-mortem damage, we soft-clipped 3 base pairs at the ends of the reads using the trimBam option in BamUtil package (v. 1.0.1466). We filtered out reads shorter than 34bp in length and under mapping quality 20. We added read groups using Picard Tools (86) before merging libraries. Finally, for samples with more than one sequenced library, we merged BAM files that had been treated independently up to this point, using the merging option in Picard.

### Population genetic analyses

We used Samtools mpileup with minimum base and mapping quality of 20 to make SNP calls and pileupCaller (https://github.com/stschiff/sequenceTools) to select one allele at random to create pseudo-haploid genotypes overlapping with the 1240k SNP panel. We filtered SNPs for linkage disequilibrium (LD) using the command --indep-pairwise (200, 25, 0.4) in PLINK1.9 (87) for *f*-statistics. We used the non-prunned dataset for remaining analyses including PCA and ADMIXTURE (Alexander et al., 2009). We used EIGENSOFT functions convertf and mergeit to merge and convert files when necessary and computed PCAs using a least squares projection of ancient samples onto modern populations using smartpca (lsqproject: YES, shrinkmode: NO). We computed *f*-statistics using the ADMIXTOOLS package (88), using qp3Pop and qpDstat. We ran *qpAdm* using different formulations to test a variety of distal and more proximal models of ancestry.

### Runs of homozygosity on pseudo-haploid data

We estimated runs of homozygosity (ROH) longer than 4 cM using hapROH (with default settings), optimised for low-coverage pseudo-haploid aDNA data (89). We restricted the analysis to individuals with at least 400,000 SNPs overlapping with the ‘1240k’ SNP panel: GOG20, GOG23, GOG24, GOG26, from medieval/Islamic contexts, and GOG50, dated to the Roman period, as well as previously published Mediterranean individuals dating to the last 2500 years (2,6–8).

### Imputation and Local Ancestry Inference (LAI) and organic homozygosity

For imputation of chromosomes 1, 11 and 21, we first generated VCF files containing all the variants in each chromosome for each individual using GATK UnifiedGenotyper (90). We combined BEAGLE 4.0 and BEAGLE 5 (91) to phase and impute the missing genotypes, as detailed in (92), making use of the 1000 Genomes (1KGP) Phase 3 reference dataset (93).

We used the VCF files to obtain LAI profiles of the three imputed chromosomes. LAI was carried out with RFMix v2 (94) with default parameters. As reference haplotypes we used the phased VCFs from the 1000 Genomes Project together with the corresponding genetic maps.

Since hapROH could not be applied to all individuals, we complemented hapROH by using the imputed chromosomes VCF files to calculate organic runs of homozygosity (oROH) and heterozygosity with in-house scripts (see Supplementary Information). We also calculated ROH statistics using PLINK (pROH) with default parameters.

## ETHICS APPROVAL AND CONSENT TO PARTICIPATE

The required permissions to access the samples were obtained with the approval of the museums involved, the Servei de Cultura i Esport de Castelló, the Direcció General de Cultura i Patrimoni, the Conselleria d’Educació, Investigació, Cultura i Esport de la Generalitat Valenciana, and the Servicio de Investigación Arqueológica Municipal del Ayuntamiento de Valencia (SIAM). Sampling was arranged to be carried out with the museum directors and/or archaeologists once permissions were granted. Samples were collected from archaeological archives in Vall d’Uixó, Gandia and Valencia.

## AVAILABILITY OF DATA AND MATERIALS

Files (FASTQ and BAM) for all newly reported individual genomes available at the European Nucleotide Archive (ENA) database under the accession number PRJEB65253.

## AUTHORS’ CONTRIBUTIONS

Study design: GOG, MBR. Field work and archaeological context: GOG, LA, MR, ARL, JPB. Laboratory work: GOG, GBF, BY, VM, CJE. Data analysis: GOG, MS, GBF, AF. Writing: GOG, MS, MBR. Supervision: MP, CJE, MBR.

Funding and other logistic support: PJ, SR, RM, FG, DGB and MBR.

## ACKNOWLEDGEMENTS

We acknowledge the people involved at the museum and local administration levels who helped throughout the bureaucratic process. Special thanks to Josep Casabó for his advice and direction during the earliest stages of sample collection. We also thank Rui Martiniano and Lara Cassidy for their advice on the bioinformatic development of this work.

## CONSENT FOR PUBLICATION

Not applicable.

## REFERENCES

1. Barceló C, Labarta A. Archivos moriscos: textos árabes de la minoría islámica valenciana 1401-1608. Valencia: Universitat de València; 2009.

2. Olalde I, Mallick S, Patterson N, Rohland N, Villalba-Mouco V, Silva M, et al. The genomic history of the Iberian Peninsula over the past 8000 years. Science. 2019;363(6432):1230–4.

3. Bycroft C, Fernández-Rozadilla C, Ruiz-Ponte C, Quintela-García I, Carracedo Á, Donnelly P, et al. Patterns of genetic differentiation and the footprints of historical migrations in the Iberian Peninsula. Nat Commun. 2018;10:551.

4. Martiniano R, Caffell A, Holst M, Hunter-Mann K, Montgomery J, Müldner G, et al. Genomic signals of migration and continuity in Britain before the Anglo-Saxons. Nat Commun. 2016;19:10326.

5. Haber M, Doumet-Serhal C, Scheib CL, Xue Y, Mikulski R, Martiniano R, et al. A Transient Pulse of Genetic Admixture from the Crusaders in the Near East Identified from Ancient Genome Sequences. Am J Hum Genet. 2019;104(5):977–84.

6. Antonio ML, Gao Z, Moots HM, Lucci M, Candilio F, Sawyer S, et al. Ancient Rome: A genetic crossroads of Europe and the Mediterranean. Science. 2019;366(6466):708–14.

7. Antonio ML, Weiß CL, Gao Z, Sawyer S, Oberreiter V, Moots HM, et al. Stable population structure in Europe since the Iron Age, despite high mobility. eLife. 2024;13:e79714.

8. Moots HM, Antonio M, Sawyer S, Spence JP, Oberreiter V, Weiß CL, et al. A genetic history of continuity and mobility in the Iron Age central Mediterranean. Nat Ecol Evol. 2023;7(9):1515–24.

9. Haak W, Lazaridis I, Patterson N, Rohland N, Mallick S, Llamas B, et al. Massive migration from the steppe was a source for Indo-European languages in Europe. Nature. 2015;

10. Olalde I, Brace S, Allentoft ME, Armit I, Kristiansen K, Booth T, et al. The Beaker phenomenon and the genomic transformation of northwest Europe. Nature. 2018;555(7695).

11. García Borja P, Blasco-Martín M, Calduch Bardoll P, Carrión P, López VC, Espinach Briansó M, et al. La inhumación tardoantigua del Hostalot-Ildum (Vilanova d’Alcolea, Castelló). Nuevas aportaciones. Quad PREHISTÒRIA Arqueol CAS℡LÓ. 2021;39:165–88.

12. Fowler C, Olalde I, Cummings V, Armit I, Büster L, Cuthbert S, et al. A high-resolution picture of kinship practices in an Early Neolithic tomb. Nature. 2022 Jan 1;601(7894):584–7.

13. Reitsema LJ, Mittnik A, Kyle B, Catalano G, Fabbri PF, Kazmi ACS, et al. The diverse genetic origins of a Classical period Greek army. Proc Natl Acad Sci. 2022;119:e2205272119.

14. Armit I, Fischer CE, Koon H, Nicholls R, Olalde I, Rohland N, et al. Kinship practices in Early Iron Age South-east Europe: Genetic and isotopic analysis of burials from the Dolge njive barrow cemetery, Dolenjska, Slovenia. Antiquity. 2023;97:403–18.

15. Silva M, Oteo-García G, Martiniano R, Guimarães J, von Tersch M, Madour A, et al. Biomolecular insights into North African-related ancestry, mobility and diet in eleventh-century Al-Andalus. Sci Rep. 2021;11(1):18121.

16. Botigué LR, Henn BM, Gravel S, Maples BK, Gignoux CR, Corona E, et al. Gene flow from North Africa contributes to differential human genetic diversity in southern europe. Proc Natl Acad Sci U S A. 2013;110(29).

17. Barral-Arca R, Pischedda S, Gómez-Carballa A, Pastoriza A, Mosquera-Miguel A, López-Soto M, et al. Meta-analysis of mitochondrial DNA variation in the Iberian Peninsula. PLoS ONE. 2016;11(7):e0159735.

18. Figueiredo Á. Death in Roman Iberia: Acculturation, resistance and diversity of beliefs and practices. Era Arqueol. 2001;ERA 3/2001.

19. Sánchez-Quinto F, Schroeder H, Ramirez O, Ávila-Arcos MC, Pybus M, Olalde I, et al. Genomic affinities of two 7,000-year-old Iberian hunter-gatherers. Curr Biol. 2012;

20. Olalde I, Allentoft ME, Sánchez-Quinto F, Santpere G, Chiang CW, DeGiorgio M, et al. Derived immune and ancestral pigmentation alleles in a 7,000-year-old Mesolithic European. Nature. 2014;507(7491):225–8.

21. Olalde I, Schroeder H, Sandoval-Velasco M, Vinner L, Lobón I, Ramirez O, et al. A Common Genetic Origin for Early Farmers from Mediterranean Cardial and Central European LBK Cultures. Mol Biol Evol. 2015;32(12):3132– 42.

22. Günther T, Valdiosera C, Malmström H, Ureña I, Rodriguez-Varela R, Sverrisdóttir ÓO, et al. Ancient genomes link early farmers from Atapuerca in Spain to modern-day Basques. Proc Natl Acad Sci U S A. 2015;112(38):11917–22.

23. Martiniano R, Cassidy LM, Ó’Maoldúin R, McLaughlin R, Silva NM, Manco L, et al. The population genomics of archaeological transition in west Iberia: Investigation of ancient substructure using imputation and haplotype-based methods. PLoS Genet. 2017;13(7).

24. Lipson M, Szécsényi-Nagy A, Mallick S, Pósa A, Stégmár B, Keerl V, et al. Parallel palaeogenomic transects reveal complex genetic history of early European farmers. Nature. 2017;551(7680):368–72.

25. González-Fortes G, Tassi F, Trucchi E, Henneberger K, Paijmans JLA, Díez-Del-Molino D, et al. A western route of prehistoric human migration from Africa into the Iberian Peninsula. Proc R Soc B Biol Sci. 2019;286(1895):20182288.

26. Valdiosera C, Günther T, Vera-Rodríguez JC, Ureña I, Iriarte E, Rodríguez-Varela R, et al. Four millennia of Iberian biomolecular prehistory illustrate the impact of prehistoric migrations at the far end of Eurasia. Proc Natl Acad Sci. 2018;

27. Villalba-Mouco V, Oliart C, Rihuete-Herrada C, Childebayeva A, Rohrlach AB, Fregeiro MI, et al. Genomic transformation and social organization during the Copper Age-Bronze Age transition in southern Iberia. Sci Adv. 2021;7(47):eabi7038.

28. Zalloua P, Collins CJ, Gosling A, Biagini SA, Costa B, Kardailsky O, et al. Ancient DNA of Phoenician remains indicates discontinuity in the settlement history of Ibiza. Sci Rep. 2018;8(1):17567.

29. Dietler M. Colonial Encounters in Iberia and the Western Mediterranean: An Exploratory Framework. In: Dietler M, Lopez-Ruiz C, editors. Colonial Encounters in Ancient Iberia: Phoenician, Greek, and Indigenous Relations. University of Chicago Press; 2009. p. 1–22.

30. Plutarch. Life of Pompey [Internet]. Parallel Lives. Available from: https://penelope.uchicago.edu/Thayer/E/Roman/Texts/Plutarch/Lives/Pompey*.html

31. Ribera i Lacomba A. La fundación de Valencia y su impacto en el paisaje. In: Historia de la Ciudad II: Territorio, Sociedad y Patrimonio, una visión arquitectónica de la historia de la ciudad de Valencia. Colegio Oficial de Arquitectos de la Comunidad Valenciana; 2002. p. 29–54.

32. Ribera i Lacomba A. El papel militar de la fundación de Valentia (138 a.C.): historia y arqueología. Defensa y territorio en Hispania de los Escipiones a Augusto: (espacios urbanos y rurales, municipales y provinciales). Coloquio celebrado en la Casa de Velázquez (19 y 20 de marzo de 2001). Casa de Velázquez; 2003. 363–390 p.

33. Ribera i Lacomba A. Depositos rituales de Valentia (Hispania). De la primera fundación republicana (138 a.C.) A la segunda augustea. In: DiGiuseppe H, Serlorenzi M, editors. I riti del costruire nelle acque violate. Roma, Palazzo Massimo: Scienze e Lettere; 2008. p. 269–94.

34. Ribera i Lacomba A. La ciudad de Valencia durante el período visigodo. In: Recópolis y la ciudad en la época visigoda. 2008. p. 303–20. (Zona Arqueológica; vol. 9).

35. Fernández Hernández G. La religión en Valencia visigoda. In: Hermosilla Pla J, editor. La ciudad de Valencia: historia, geografía y arte de la ciudad de Valencia [Internet]. 2009. p. 150–3. (Historia; vol. 1). Available from: https://roderic.uv.es/rest/api/core/bitstreams/8e74c070-63af-4ed6-8c6a-30937d4c3772/content

36. Coscollá Sanz V. La Valencia musulmana. Carena Editors; 2003.

37. Rosenwein BH. Umayyad diplomacy: The Treaty of Tudmir (713). In: Reading the Middle Ages: Sources from Europe, Byzantium, and the Islamic World (Third Edition). Broadview; 2018. p. 78.

38. König DG. 713: The Treaty of Tudmir as a Testimony to the Muslim Subjection of the Iberian Peninsula. Transmediterranean Hist. 2020;2(1).

39. Ribera A, Rosselló M, Macias JM. Historia y arqueología de dos ciudades en los siglos VI-VIII d. C. Valentia y València la Vella. Antigüedad Crist. 2020;37:63–106.

40. Macias JM, Ribera A, Rosselló M, Rodríguez F, Caldés Ò. València la Vella: A Visigothic city to place in history? Post -Class Archaeol. 2023;13:31–54.

41. Watt WM, Cachia P. A History of Islamic Spain. Edinburgh University Press; 1965. 210 p.

42. Haliczer S. Inquisition and Society in the Kingdom of Valencia, 1478– 1834. University of California Press; 1990.

43. Baydal V, Esquilache F. Exploitation and differentiation: economic and social stratification in the rural Muslim communities of the Kingdom of Valencia, 13th–16th centuries. In: Beyond lords and peasants: rural elites and economic differentiation in pre-modern Europe. Universitat de Valencia; 2014. p. 37–67.

44. Alexander MM, Gerrard CM, Gutiérrez A, Millard AR. Diet, society, and economy in late medieval Spain: Stable isotope evidence from Muslims and Christians from Gandía, Valencia. Am J Phys Anthropol. 2015;156(2):263–73.

45. Harvey LP. Muslims in Spain, 1500 to 1614. University of Chicago Press; 2005.

46. Burns RI. Islam under the Crusaders: Colonial Survival in the Thirteenth-Century Kingdom of Valencia. Islam under the Crusaders: Colonial Survival in the Thirteenth-Century Kingdom of Valencia. Princeton University Press; 1973.

47. Burns RI. Medieval Colonialism: Postcrusade Exploitation of Islamic Valencia. Medieval Colonialism: Postcrusade Exploitation of Islamic Valencia. Princeton University Press; 1975.

48. Lapeyre H. Geografía de la España morisca. Universitat de València; 2011.

49. Jónsson M. The expulsion of the Moriscos from Spain in 1609–1614: The destruction of an Islamic periphery. J Glob Hist. 2007;2(2):195–212.

50. Carr M. Blood and Faith: The Purging of Muslim Spain 1492-1614. C. Hurst and Co.; 2009.

51. Alexander MM, Gutiérrez A, Millard AR, Richards MP, Gerrard CM. Economic and socio-cultural consequences of changing political rule on human and faunal diets in medieval Valencia (c. fifth–fifteenth century AD) as evidenced by stable isotopes. Archaeol Anthropol Sci. 2019;11:3875–93.

52. Oteo-García G. Archaeogenetics of Southwest Europe [Internet] [PhD Thesis]. [Huddersfield]: University of Huddersfield; 2020. Available from: https://eprints.hud.ac.uk/id/eprint/35459

53. Mallick S, Reich D. The Allen Ancient DNA Resource (AADR): A curated compendium of ancient human genomes. 2023.

54. Derenko M, Malyarchuk B, Grzybowski T, Denisova G, Rogalla U, Perkova M, et al. Origin and post-glacial dispersal of mitochondrial DNA haplogroups C and D in northern Asia. PloS One. 2010;5(12):e15214.

55. García Prósper E. Los ritos funerarios de la necrópolis romana de la calle Quart de Valencia (Siglos II a.C-III d.C) [Tesis doctoral]. [Valencia, España]: Universitat de València; 2016.

56. Oteo-García G, Alapont-Martín L, Pascual Beneyto J, Foody MGB, Yau B, Pala M, et al. Late Roman tombs at Sanxo Llop (Gandía, Valencia): Exogamy and kinship in a particular funerary structure. In: Death and the Societies of Late Antiquity: New methods, new questions? (Archéologies méditerranéennes). Aix-en-Provence: Presses universitaires de Provence; 2023. p. 119–27.

57. Olivé-Busom J, López-Costas O, Márquez-Grant N, Kirchner H. Estudio antropológico de las alquerías de Benizahat y Zeneta (Vall d’Uixó, Castellón). Una ventana a la vida rural andalusí. SAGVNTVM PLAV. 2021;53:193–212.

58. Agranat-Tamir L, Waldman S, Martin MAS, Gokhman D, Mishol N, Eshel T, et al. The Genomic History of the Bronze Age Southern Levant. Cell. 2020;181:1146–57.

59. Lazaridis I, Alpaslan-Roodenberg S, Acar A, Açıkkol A, Agelarakis A, Aghikyan L, et al. The genetic history of the Southern Arc: A bridge between West Asia and Europe. Science. 2022;377(6609):eabm4247.

60. Solé-Morata N, Villaescusa P, García-Fernández C, Font-Porterias N, Illescas MJ, Valverde L, et al. Analysis of the R1b-DF27 haplogroup shows that a large fraction of Iberian Y-chromosome lineages originated recently in situ. Sci Rep. 2017;7(1):7341.

61. Soares P, Ermini L, Thomson N, Mormina M, Rito T, Röhl A, et al. Correcting for Purifying Selection: An Improved Human Mitochondrial Molecular Clock. Am J Hum Genet. 2009;84(6).

62. Lee J, Miller BK, Bayarsaikhan J, Johannesson E, Ventresca Miller A, Warinner C, et al. Genetic population structure of the Xiongnu Empire at imperial and local scales. Sci Adv. 2023;9(15):eadf3904.

63. Jeong C, Wang K, Wilkin S, Taylor WTT, Miller BK, Bemmann JH, et al. A Dynamic 6,000-Year Genetic History of Eurasia’s Eastern Steppe. Cell. 2020;183(4):890–904.e29.

64. Heather PJ. Empires and Barbarians. Macmillan; 2009.

65. Arauna LR, Mendoza-Revilla J, Mas-Sandoval A, Izaabel H, Bekada A, Benhamamouch S, et al. Recent Historical Migrations Have Shaped the Gene Pool of Arabs and Berbers in North Africa. Mol Biol Evol. 2017;34(2):318–29.

66. Forbes JD. Africans and Native Americans: The Language of Race and the Evolution of Red-Black Peoples. University of Illinois Press; 1993. 145 p.

67. Al-Rasheed M, Vitalis R. Counter-Narratives: History, Contemporary Society, and Politics in Saudi Arabia and Yemen. Palgrave Macmillan US; 2004. 136 p.

68. Bernards M, Nawas JA. Patronate And Patronage in Early And Classical Islam. BRILL; 2005. 219 p.

69. Ubieto Arteta A. Orígenes del reino de Valencia: cuestiones cronológicas sobre su reconquista. Zaragoza: Anubar; 1975.

70. Flores-Bello A, Bauduer F, Salaberria J, Oyharçabal B, Calafell F, Bertranpetit J, et al. Genetic origins, singularity, and heterogeneity of Basques. Curr Biol. 2021;31(10):2167–2177.e4.

71. Martín Corrales E. Chapter 2: The Spain That Enslaves and Expels: Moriscos and Muslim Captives (1492 to 1767–1791). In: López-Morillas C, editor. Muslims in Spain, 1492-1814. Brill; 2020. p. 67–94.

72. Cortés Alonso V. Procedencia de los esclavos negros en Valencia (1482-1516). Rev Esp Antropol Am. 1972;7(1):123–52.

73. Marzal Palacios FJ. Una presencia constante: los esclavos sarracenos en Valencia (siglos XIII-XVI). Sharq Al-Andal. 2002;16–17.

74. Marzal Palacios FJ. La esclavitud en Valencia durante la Baja Edad Media (1375-1425). PhD thesis. Universitat de Valencia; 2006.

75. Chacón-Duque JC, Adhikari K, Fuentes-Guajardo M, Mendoza-Revilla J, Acuña-Alonzo V, Barquera R, et al. Latin Americans show wide-spread Converso ancestry and imprint of local Native ancestry on physical appearance. Nat Commun. 2018;9(1):5388.

76. Yang DY, Eng B, Waye JS, Dudar JC, Saunders SR. Improved DNA extraction from ancient bones using silica based spin columns. Am J Phys Anthropol. 1998;105(4):539–43.

77. MacHugh D, Edwards C, Bailey J, Bancroft D, Bradley D. The extraction and analysis of ancient DNA from bone and teeth: a survey of current methodologies. Anc Biomol. 2000;3:81–102.

78. Rohland N, Hofreiter M. Ancient dna extraction from bones and teeth. Nat Protoc. 2007;

79. Meyer M, Kircher M, Gansauge MT, Li H, Racimo F, Mallick S, et al. A high-coverage genome sequence from an archaic Denisovan individual. Science. 2012;338(6104):222–6.

80. Gamba C, Hanghøj K, Gaunitz C, Alfarhan AH, Alquraishi SA, Al-Rasheid KAS, et al. Comparing the Performance of Three Ancient DNA Extraction Methods for High-Throughput Sequencing. Mol Ecol Resour. 2016;16(2):459– 69.

81. Cassidy LM, Martiniano R, Murphy EM, Teasdale MD, Mallory J, Hartwell B, et al. Neolithic and Bronze Age migration to Ireland and establishment of the insular Atlantic genome. Proc Natl Acad Sci. 2016;113(2):368–73.

82. Andrews S. FASTQC A Quality Control tool for High Throughput Sequence Data. Babraham Inst. 2015;

83. Schubert M, Ginolhac A, Lindgreen S, Thompson JF, AL-Rasheid KAS, Willerslev E, et al. Improving ancient DNA read mapping against modern reference genomes. BMC Genomics. 2012;13:178.

84. García-Alcalde F, Okonechnikov K, Carbonell J, Cruz LM, Götz S, Tarazona S, et al. Qualimap: evaluating next-generation sequencing alignment data. Bioinformatics. 2012;28(20):2678–9.

85. Li H, Handsaker B, Wysoker A, Fennell T, Ruan J, Homer N, et al. The Sequence Alignment/Map format and SAMtools. Bioinformatics. 2009;25(16):2078–9.

86. Broad Institute. Picard Toolkit [Internet]. 2019. Available from: http://broadinstitute.github.io/picard/

87. Purcell S, Neale B, Todd-Brown K, Thomas L, Ferreira MAR, Bender D, et al. PLINK: A tool set for whole-genome association and population-based linkage analyses. Am J Hum Genet. 2007;81(3):559–75.

88. Patterson N, Moorjani P, Luo Y, Mallick S, Rohland N, Zhan Y, et al. Ancient admixture in human history. Genetics. 2012;192:1065–93.

89. Ringbauer H, Novembre J, Steinrücken M. Parental relatedness through time revealed by runs of homozygosity in ancient DNA. Nat Commun. 2021;12:5425.

90. McKenna A, Hanna M, Banks E, Sivachenko A, Cibulskis K, Kernytsky A, et al. The genome analysis toolkit: A MapReduce framework for analyzing next-generation DNA sequencing data. Genome Res. 2010;20(9):1297–303.

91. Browning SR, Browning BL. Rapid and accurate haplotype phasing and missing-data inference for whole-genome association studies by use of localized haplotype clustering. Am J Hum Genet. 2007;81(5):1084–97.

92. Hui R, D’Atanasio E, Cassidy LM, Scheib CL, Kivisild T. Evaluating genotype imputation pipeline for ultra-low coverage ancient genomes. Sci Rep. 2020;10(1):18542.

93. Auton A, Abecasis GR, Altshuler DM, Durbin RM, Bentley DR, Chakravarti A, et al. A global reference for human genetic variation. Nature. 2015;526(7571):68–74.

94. Maples BK, Gravel S, Kenny EE, Bustamante CD. RFMix: a discriminative modeling approach for rapid and robust local-ancestry inference. Am J Hum Genet. 2013;93(2):278–88.

