## Supplementary Information for "Medieval genomes from eastern Iberia illuminate the role of Morisco mass deportations in dismantling a long-standing genetic bridge with North Africa"

Table of contents:


- Supplementary Material and Methods … 2
- Supplementary Tables list ………………... 15
- Supplementary Figures ……………………. 17
- Supplementary References ……………… 28


### Supplementary Materials and Methods

#### Sampling locations

Once permissions to access the samples were obtained, sampling for the following locations was arranged to be carried out with the museum directors or archaeologists. Once the samples have been screened for the content of endogenous DNA, the best preserved samples were selected for further sequencing. These twelve individuals were drawn as the best from the total 35 screened samples which had yielded endogenous DNA proportions ranging from 0.01% to 41.58%. The average endogenous DNA content was about 7%. The sequencing of the DNA recovered from bones that included petrouses, molars and a metatarsus yielded average nuclear coverages, after mapping quality filters, ranging from 0.21× to 2.34×.

• Necropolis de La Union (Vall d’Uixó): cemetery from the Visigothic period with over 40 inhumations (39.816157, -0.234099). This site is peculiar since almost all the graves had several individuals buried one on top of another. The archaeological interpretation is that the individuals in shared graves share some familial link. They all appear to be victims of a violent event, and they died within a short period of time. Many individuals display signs of violence as revealed by anthropological study. We sampled five individuals from this site but none yielded enough endogenous DNA contents to be sequenced further. No radiocarbon dates available. Samples: GOG15, GOG16, GOG17, GOG18, GOG19 (1).

• Islamic Maqbaras (Vall d'Uixó): these are a series of burials discovered during various excavations in the 1990s across the Vall d'Uixó. They have been identified as the islamic cemeteries of the agricultural hamlets or farms (Alquería Benigafull (39.823088, -0.235724), Alquería Benizahat (39.824500, -0.229082), and Alquería Ceneja (39.822567, -0.23749) that formed the primitive cores of the Vall d'Uixo settlement. The locations of Alquería Ceneja and Alquería Benifagull were on the high ground of the modern town. The Alquería Benizahat was on the lower part of the modern town. These settlements eventually fused to originate the modern municipality (2). Archaeological inferences estimated that the settlements on the high ground are older than the hamlets in the lower ground. We sampled 9 individuals from four of these sites. The samples were initially believed to be from between the 11th and 14th centuries. However, the radiocarbon dating performed on three of the samples revealed a wider time frame, from the 8th to the 14th century. Samples: GOG20, GOG21, GOG22, GOG23, GOG24, GOG25, GOG26, GOG29, GOG30 (1).

• Sanxo Llop and La Vital (Gandia): two excavation sites in very close proximity to one another. La Vital (38.969286, -0.167254) is an archeological site with evidence of human occupation going back to the Late Neolithic. From this location we collected three samples from an early medieval pre-Islamic context, however endogenous DNA contents were insufficient for further sequencing and these samples were not included in further analyses. In Sanxo Llop (38.963488, -0.169599), excavations revealed a burial ground dating to late Antiquity, overlapping between the late Visigothic period and early Islamic decades, at least between 660-800 CE according to available radiocarbon dates. We sampled two individuals (an adult and a sub-adult) that revealed good preservation of endogenous DNA. Previously available radiocarbon dates indicate the shared burial dates to the period between 6th-7th centuries CE. The two individuals were deposited in a round pit that had likely been used for other purposes in the past. Samples: GOG31, GOG32, GOG33, GOG34, GOG35 (1,3).

• Cementerio de San Lorenzo (Valencia): Chrisitian cemetery inside the walled perimeter of medieval Valencia (39.477944, -0.375865). In use since at least the 15th century up until the mid 19th century. In the 19th century the growing consensus that cemeteries inside cities were a health hazard led to the development of burial grounds outside urban areas. In 1841 following the confiscation of urban cemeteries, all the known and recent graves were moved to the new general cemetery of Valencia. Excavations in 1999 on the site of the former cemetery at Plaza Cisneros found the older non-translocated medieval graves, spanning from the 14th century to the 17th. The same area covered by the medieval cemetery had been used in Late Roman times for burials and the excavation found the remains of two of these Roman tombs. We sampled 8 individuals from the 14-17th century period and the two individuals from the Roman graves. The remains from the Roman burials were poorly preserved but one yielded exceptionally high amounts of endogenous

DNA. Samples: GOG49, GOG50, GOG53, GOG54, GOG55, GOG56, GOG57, GOG58, GOG59, GOG60 (1).

• Other locations: we sourced other bone material that was unsuccessful in the recovery of ancient DNA. These other samples were scattered an include one Islamic burial (GOG45) from an excavation in Calle Ripalda (Valencia), three Islamic samples (GOG13, GOG14, GOG46) from Plaza del Almudín in Segorbe (a site previously studied in (4)) and two medieval samples (GOG51, GOG52) from Monasterio San Vicente de la Roqueta (Valencia) (1).

**Processing of ancient bones**

The handling of ancient DNA material was carried out at the dedicated Ancient DNA Facility run by the Archaeogenetics Research Group at the University of Huddersfield. The Ancient DNA Facility is isolated from other molecular biology facilities handling post-PCR products or modern sources of DNA since it is located in the Technology Building of the Queensgate Campus (Huddersfield). In every work session, handlers were outfitted with a full-body Tyvek suit, a hairnet and face mask and a double layer of gloves at all stages. The facility includes two independent rooms dedicated to sample drilling and DNA extraction/library-preparation respectively. The protocol required all materials and surfaces to be regularly cleaned with LookOut® DNA Erase (Sigma-Aldrich), bleach and by exposure to UV-radiation after each session. To reduce risk of bacterial and other contamination, the samples selected to be processed by drilling were UV-radiated for a total of 60 minutes (30 minutes for each side) before being carried to the sample drilling room. We cleaned the sampling surfaces of the petrous bones or molars by air-abrasion, with 29 µm aluminium oxide powder (OEA Labs) and a SWAM-Blaster® compressed air abrasive system (Crystal Mark). We used a Micromotor System Maxima drill with a 22 mm diameter diamond cutting edge for sampling. For teeth we separated the root and crown to powder the roots for DNA extraction. For petrous bones, we extracted a bone wedge from the densest part (5). In the case of other bones (phalanx, tarsus) we targeted the epiphysis. Once a piece was obtained, we powdered the samples using a Mixer Mill (Retsch MM400) for 45 seconds at a frequency of 30 Hz/s.

**Radiocarbon dating**

We dated petrous bones from four individuals (GOG20, GOG23 and GOG26 from La Vall d'Uixo, and GOG50 from the Cementerio San Lorenzo) at 14Chrono, the radiocarbon dating lab at Queen's University (Belfast). We calibrated the resulting dates in cal. CE using OxCal version 4.4.4 and the most recent calibration curve, IntCal20 (6,7). Sample GOG34 had been dated in the past at Beta Analytic (Miami) by means of AMS measurements made with NEC SSAMS accelerator mass spectrometers (SI).

**Extraction of ancient DNA**

We followed a modified protocol extraction of DNA from ancient remains (8–10), and used Fisher reagents. The extraction buffer (EB) contained 20mM of Tris HCL, pH 8; 50mM of EDTA, pH 8, RNase and Proteinase free, and 0.5% of SDS (DNase, RNase and protease free, heated to a temperature of 37ºC). All components were exposed to UV-light for 15 minutes before addition of proteinase K. Once ready, 1 mL of extraction buffer was added to the tube with the powder, which were then incubated rotating for approximately 24 hours at 37 ºC. Following incubation, the tubes were centrifuged at 13,000 rpm for 15 minutes to separate mineral particles from the supernatant (which was retained at -20 C). We added 1 ml of EB to the remaining pellet, vortexed the tubes to resuspend it. and repeated the  incubation for another 24 hours rotating at 37ºC. After the second incubation the tubes were centrifuged at 13,000 rpm again for another 15 minutes and the . supernatant was transferred to 6 mL Corning® Spin-X® UF Concentrator tubes. To these new tubes a quantity of 3mL of 10mM Tris HCL (pH 8) was added and then centrifuged for 20 minutes at 2,500 rpm twice ( the flow-through was discarded after the first centrifugation, and 3mL of 10mM Tris HCL (pH 8) added again).  We retained a final volume of  ~100 µL, which was  transferred to new silica columns (MinElute® PCR Purification Kit, commercialised by Qiagen) for purification following standard protocol by the manufacturer, plus addition of 0.05% Tween-20 (0.03µL per sample) to 59.97 µL per sample of EB Buffer to reduce absorption of DNA to plastics and keep viability of extraction in the long run. The final 100 µL volume of DNA extracted was at 4ºC. Finally, a small volume of each DNA extraction was taken to quantify with a QubitTM 3.0 Fluorometer (ThermoFisher Scientific), using the Qubit® dsDNA HS Assay Kit (Invitrogen).

**Library preparation and sequencing**

The protocol from Meyer *et al.* (11), with modifications introduced in Gamba *et al.* and Cassidy *et al.* (12,13), was followed to make the necessary library preparations for the samples. The initial step  of DNA fragmentation was skipped since ancient DNA by nature is already highly fragmented. In between all main stages (i, ii, iii, iv and v) of library preparation, clean-up steps were performed using the MinElute PCR Purification Kit, according to manufacturer instructions, and adding Tween 0.05% to EB Buffer to obtain EBT Buffer in the same way as detailed above.

After a initial screening sequencing round using a Illumina MiSeq platform (Trinity Genome Sequencing Laboratory, Trinity College Dublin, Ireland), the libraries  with sufficient endogenous DNA for further sequencing were UDG-treated  before library-preparation as follows: addition of 5.0 µL of USER® enzyme (Uracil-Specific Excision Reagent by New England BioLabs®) to 16.5µL of DNA extract and incubated for 3 hours at 37ºC in order to remove uracil residues derived from post-mortem damage characteristic of ancient DNA (14–17) .

The library preparation protocol consisted of various stages which included blunt-end repair (i), adapter ligation (ii), followed by an adapter fill-in reaction with Bst DNA polymerase (iii). Indexing oligo sequences were added by amplification with IS4 primer (iv). Finally libraries to be sequenced together in the same lane were pooled together (v).

Step I) For each sample, we merged the cleaned DNA volume of 21.5µL resulting from the USER-treatment, together with 3.5µL of NEBNext End Prep Enzyme Mix, 7µL of 10X of NEBNext End Repair Reaction Buffer (both included in the NEBNext® End Repair Module, New England BioLabs®), and 38µL of ddH2O (sterile ultrapure water). The final volume (for one sample) was 70.0µL that was later incubated at 25ºC for 15 minutes, followed by 5 minutes at 12ºC, and purified with the MinElute PCR Purification Kit.

Step II): The adapter mixes of P5 and P7 (20µM each) (by Sigma-Aldrich) were pooled together with 1µL of T4 DNA ligase I (5U/µL), 10µL of ddH2O (sterile ultrapure water), 4µL of 10X 101T4 DNA ligase buffer by Thermo Scientific , and 4µL 50% PEG-4000 (Thermo Scientific). The final volume was 20µL per sample and was pooled together with another 20 µL of eluate from the previous Step I, then it was incubated at 22ºC for 30 minutes. In this step, adapters were ligated by the activity of T4 DNA ligase catalysis of phosphodiester bonds between 5’ and 3’-ends in dsDNA.

Step III) For the adapter fill-in step, we merged the 20µL of DNA from Step II with 13.5µL of ddH2O (sterile ultrapure water), 4µL ThermoPol® Reaction Buffer 10X, 1µL of dNTP (10mM each), and 1.5µL of Bst DNA polymerase (Large Fragment, 8U/µL), resulting in a total volume of 40 µL to be incubated for 30 minutes at 37ºC, followed by an extra 20 minutes at 80 ºC necessary to inactivate the Bst DNA polymerase and terminate the reaction.

Step IV) Library amplification reactions were prepared in the Ancient DNA lab and the final amplification step was carried out in the post-PCR lab space The reaction consisted of 41 µL of Accuprime Pfx SuperMix (Thermo Scientific), 1µL of primer IS4 (10µM), 2µL of appropriate indexing oligo (both made by Sigma-Aldrich) plus 6µL of sample library from Step III. Total reaction volume is 50µL. The PCR reaction protocol consisted of an initial denaturation phase at 95ºC for 5 minutes, followed by 12 cycles of denaturation at 95ºC for 15 seconds, annealing at 60ºC for 30 seconds, extension at 68ºC for 30 seconds, and a final extension at 68ºC for 5 minutes. Finally, we performed the last clean-up step with the MinElute PCR Purification Kit.

Step V) We measured the concentration of the libraries with a QubitTM 3.0 Fluorometer, using the Qubit® dsDNA HS Assay Kit and checked the fragment size distribution with a Bioanalyzer (Agilent), using the Agilent High Sensitivity DNA Kit. Libraries were evaluated visually and if deemed successful (majority of fragment lengths in the libraries peaking at 150-200 bp), libraries were pooled together. We sent the libraries to Macrogen (Seoul, South Korea) for whole genome next-generation sequencing (NGS) in Illumina HiSeq 4000 platforms.

### Processing Next-Generation Sequencing data

The paired-end raw FASTQ files were evaluated using FastQC (version 0.11.7 by Babraham Bioinformatics) to check for quality of the high throughput sequence data. Results of the checks were inspected visually. Once we cleared the FASTQ viability we proceeded to remove adapter sequences and merged paired-end files. We used leeHom (18) to merge paired-end FASTQ files without and to remove adapters in one step by making use of its algorithm, with the flag –ancientdna.

### Mapping NGS data

We used Burrows Wheeler Aligner (BWA) with the commands samse and aln to generate BAM files mapped to the Human Reference Build 37 (hg19/GRCh37.p13). Ancient DNA specifications ) were used to disable the minimum seed length (-l option) and allow the aligning algorithm to be more flexible and map more reads increasing coverage (-n 0.01, -o 2), as well as applying map quality filtering 20 . We used Qualimap (19) to retrieve alignment metrics as quality control.

We removed duplicates from the BAM files  with Samtools rmdup. Once the duplicates were removed, quality filters were applied. A minimum mapping quality of 20 was chosen to reduce reference bias than the also commonly used threshold of 30. Minimum read lengths were set at 34 base pairs long (13,20). PMD patterns were evaluated visually by looking at the plots generated with mapDamage (v.2.0.7). We soft-clipped three base pairs at the end of each read using the trimBam (–clip) function in bamUtil to remove the vast majority of the deamination damage, given that the sequences are already uracil–DNA–glycosylase (UDG) treated (21).

We added read groups at library-level using Picard Tools, and we merged all libraries from the same sample (which had been treated independently until now) into one file using the merging option in Picard.

**Data authenticity**

Anti-contamination measures were in place while drilling, extracting aDNA and during library preparation in the Ancient DNA Facility as explained above. Negative controls were also introduced in the chain leading to library preparation and sequencing with Illumina MiSeq (Trinity Genome Sequencing Laboratory, Trinity College Dublin, Ireland) in the form of air (empty tube opened for some time before sample drilling began) and water control (2ml of ddH2O shaken in the sample mixer used to powder bone, after bleaching and UV exposure). The DNA quantification and sequencing results of these controls showed that the levels of contamination by exogenous DNA were negligible. We further checked authenticity of the data by checking the patterns of post-mortem damage and DNA fragmentation in the GOG samples with MapDamage v.2.0.7  and BamDamage (22,23). We checked contamination estimates on mtDNA with Schmutzi. All non-USER-treated libraries generated for screening samples presented the typical misincorporation patterns of ancient DNA, which were also detectable in the non-treated screening libraries and in very low levels in the USER-treated libraries (Figure S2). We also checked levels of mtDNA contamination in the samples with Schmutzi. We further evaluated that the mitochondrial haplotypes of each individual were consistent with one haplogroup only.

**Biological sex determination**

Genetic sex determination was established with Ry score (24) on all individual libraries before and after merging libraries.

**Classification of uniparental markers**

Mitochondrial reads were mapped to the revised Cambridge Reference Sequence (rCRS) mitochondrial genome (NC_012920.1). We obtained the mitochondrial mutations using GATK (v.3.7-0) HaplotypeCaller (25) and we further evaluated the haplotypes on IGV v.2.3 (26) to visually inspect any heteroplasmic positions. Haplogroup classification was made using HaploGrep 2.0 (27) following the nomenclature in PhyloTree (Build 17, February 2016) (28).

Haplotypes from male samples with Y chromosome data were classified following with the 2019 ISOGG list of mutations (International Society of Genetic Genealogy) using Yleaf (29) For samples whose Yleaf classification was doubtful we double checked using pathPhynder (30) guide tree a tool that integrates ancient and modern variation taking into account all informative Y-chromosome markers.

**Kinship determination**

To infer kinship relationships up to second degree we used the software Relationship Estimation from Ancient DNA (READ) (31). READ is optimised to manage low-coverage pseudo-diploid data in EIGENSTRAT format. It tests systematically pairs of individuals in a given dataset and is able to classify the type of relatedness degree as non-related, second degree (e.g. grandparent-grandchild, half-siblings, uncle/aunt, nephew/niece), first degree (parent-offspring, siblings), and identical (twins or duplicated individual). All combinations of samples, with the same SNP list as for the other analyses, were attempted regardless of location and time period to confirm no cross-contamination happened between samples which would show up as unexpected artificial relatedness. Only samples GOG34 and GOG35 were found to be biological relatives (1st degree).

**SNP calling from BAM files**

To call the variants we used two lists of SNPs: I) ~600k list used in the Human Origins dataset (Lazaridis et al., 2014) and II) ~1.2M SNPs included in the ‘1240k’ targeted enrichment protocol. The SNPs were called on all ancient samples using a combination of SAMtools mpileup and SequenceTools pileupCaller (with quality filters q20, Q20 enabled) (https://github.com/stschiff/sequenceTools). Pseudo-haploid genotypes were called by randomly choosing one allele from each site where there was read coverage, using pileupCaller.

**Data merging with public datasets**

In total, 12 individual genomes passed endogenous preservation and quality control thresholds.

We merged the newly generated ancient genomes reported here with genotypes from relevant ancient and present-day populations from across the Mediterranean and neighbouring regions included in the Allen Ancient DNA dataset (32) We made use of modern populations from the dataset to compute the PCA onto where ancient individuals were projected. We avoided comparisons with capture generated samples (which limited available sources from Iberia) to limit biases in f-statistics tests and qpAdm.

We also merged the dataset with the North African dataset from (33) for comparisons in f-statistics and expanding the diversity of North African groups.

For haplotype-based analyses we relied on the 1000 Genomes Project publicly available data.

**Simulation of hybrid genomes**

We simulated hybrid genotypes imitating the offspring between two different parental sources of Maghrebi and Iberian origin. The simulation follows Mendelian segregation rules for a case of one gene with two alleles but does not take into account recombination. If both parents are homozygous for the reference or alternative allele then the hybrid offspring will carry two reference or alternate alleles respectively. If one parent is homozygous for allele A, and the other for allele B, then the offspring will always be assigned a heterozygous genotype. In situations where there could be more than one resulting genotype, a function assigned the genotype randomly according to the expected probabilities. We then projected these hybrids onto a PCA of a merged dataset with ~20k SNPs that included modern individuals from the HO dataset and Arauna et al.,dataset.

**Clustering Analyses**

For Principal Component Analysis I used the EIGENSOFT tool smartpca (34). The PCA dimensionality reduction was carried out using a subset of modern populations from the ~600k SNPs Human Origins dataset (35). The subset of modern populations included all individuals available from Europe, the Caucasus, Iran, the Near East, Arabia and North Africa. The 434 public ancient genomes from Spain, Anatolia, the Fertile Crescent and Morocco were later projected onto the PCA built with the modern populations (option lsqproject: YES). Outliers were not excluded.

We used the model-based clustering approach of ADMIXTURE (36) to estimate ancestry components in the newly generated samples, together with the same ancient populations included in the PCA. We applied a filter for linkage disequilibrium in PLINK. (parameters 200, 25 and 0.4). This filter decreased the final number of SNPs used in ADMIXTURE to about ~200k SNPs. We ran ADMIXTURE in supervised and unsupervised mode (Figures 3A, S6). In unsupervised mode we only included ancient populations from K=2 to K=10 with the cross-validation flag activated (–cv). The lowest median CV error before a steep rise was for K=4.

**F-Statistics**

Formal tests to evaluate treeness, gene flow and admixture were performed with *f*-statistics (37–39) included in the AdmixTools package by the Harvard Reich Lab ((40). Default parameters were used. The f3 tests were used both in the form of outgroup-f3 and admixture-f3 depending on the scenario to be investigated.

We used *qpWave/qpAdm* to test a variety of distal and more proximal models of ancestry. We started by applying a modified version of a distal model optimised for post-Bronze Age West European populations first described in (41), using *WHG.SG*, *Balkan_N.SG* and *Yamnaya.SG* as fixed sources to represent Western European Hunter-Gatherer-, Early European Farmer- and Steppe-related ancestries, respectively. Following the observation that the model is rejected for most individuals (Table S14), we repeated the model adding Morocco_EN.SG to the source populations, as a proxy for North African-related ancestry.

To test more temporally proximal sources we ran 1-source *qpWave* models, rotating through a reference list including outgroups and historical

populations: *South_Africa_400BP.SG*, *Russia_Yana_UP.SG*, *Japan_Honshu_EarlyJomon.SG*, *Brazil_Sumidouro.SG*, *Morocco_EN.SG*, *Italy_RomanImperial.SG*, *Lebanon_Roman.SG*, *CanaryIslands_Guanche.SG*, *Portugal_LateRoman.SG* *(Portugal_Miroico_LateRoman.SG+Portugal_MonteDaNora_LateRoman.SG)*, *England_IA_Roman.SG* (https://github.com/pontussk/qpAdm_wrapper).
For individuals with no single-source *qpWave* model accepted or with more than one model accepted, we tested all possible combinations of two sources, using the same set of references.
We restricted the dataset to data generated using whole-genome shotgun sequencing to avoid potential biases with capture generated samples (which limited the available sources from Iberia) to avoid potential biases in f-statistics tests and *qpAdm*.

**Analysis of imputed genomes**

For LAI, ROH and heterozygosity analyses we made use of the imputed VCF files of the chromosomes. LAI was carried out with RFMix v2 (42). As reference haplotypes we used the phased VCFs from the 1000 Genomes Project together with the corresponding genetic maps.
We obtained information about ROH in different manners. One involved using hapROH. Another PLINK (with parameters --homozyg --homozyg-density 50 --homozyg-gap 100 --homozyg-kb 500 --homozyg-snp 50 --homozyg-window-het 1 --homozyg-window-snp 50 --homozyg-window-threshold 0.05) which only identifies certain segments of the chromosome as a ROH based on the stringency of the parameters given. The other manner we used was recording the length of each segment in between every single heterozygous position along the full extent of each chromosome. We call these segments, “organic

ROH” (oROH) as opposed to the “PLINK ROH” since they are identified free of prior assumptions.

Sliding heterozygosity through the chromosomes was computed after filtering and retaining transitions and transversions from the 1000G Phase 3 list. We then divided his genome into 500 marker windows and calculated the percentage of heterozygous sites within each window, using an overlapping step of size 250 (43). We then used this information to obtain a value of entropy (H’) following the formula of the Shannon Diversity Index.

**Supplementary Tables list:**

- Table S1: Details of final sequencing results for the ancient samples studied.
- Table S2:  Details of all samples screened for endogenous content.
- Table S3: Details and about the new radiocarbon dates generated.
- Table S4: Homozygous segments identified in the imputed chromosomes using PLINK with default parameters.
- Table S5: Total number of RoH, total length of ROH measured in Kb, and average length of ROH in Kb. Includes the newly generated ancient genomes as well as the entire 1000 Genomes dataset.
- Table S6a: Measurements by sliding windows of number of variant sites in imputed chromosome 1 for the ancient genomes reported in this study. Used in Figure S8.
- Table S6b: Measurements by sliding windows of number of variant sites in imputed chromosome 11 for the ancient genomes reported in this study. Used in Figure S8.

- Table S6c: Measurements by sliding windows of number of variant sites in imputed chromosome 21 for the ancient genomes reported in this study. Used in Figure S8.
- Table S7a: Cumulative ROH curve for chromosome 1. Used in Figure S9.
- Table S7b: Cumulative ROH curve for chromosome 11. Used in Figure S9.
- Table S7c: Cumulative ROH curve for chromosome 21. Used in Figure S9.
- Table S8a: Organic ROH fragments in chromosome 1 identified in the ancient genomes reported in this study.
- Table S8b:  Organic ROH fragments in chromosome 11 identified in the ancient genomes reported in this study.
- Table S8c:  Organic ROH fragments in chromosome 21 identified in the ancient genomes reported in this study.
- Table S9: Summary statistics of organic ROH used for Figure 4B.
- Table S10: Shannon entropies for all imputed chromosomes in the ancient individuals reported.
- Table S11: Outgroup-f3 test with the medieval ancient genomes and modern populations merged overlapping the Allen Ancient DNA Resource (AADR 1240k) and Arauna et al., (2017) datasets. Used in Figure 2B.
- Table S12: Outgroup-f3 test for sample GOG50 with modern and ancient populations.
- Table S13: qpAdm proximal 1 source and 2 source models (rotating).
- Table S14: qpAdm distal models with 3 and 4 sources (fixed).
- Table S15: Genetic sex identification.
- Table S16: Haplogroup classification of Y chromosomes.

**Supplementary Figures**

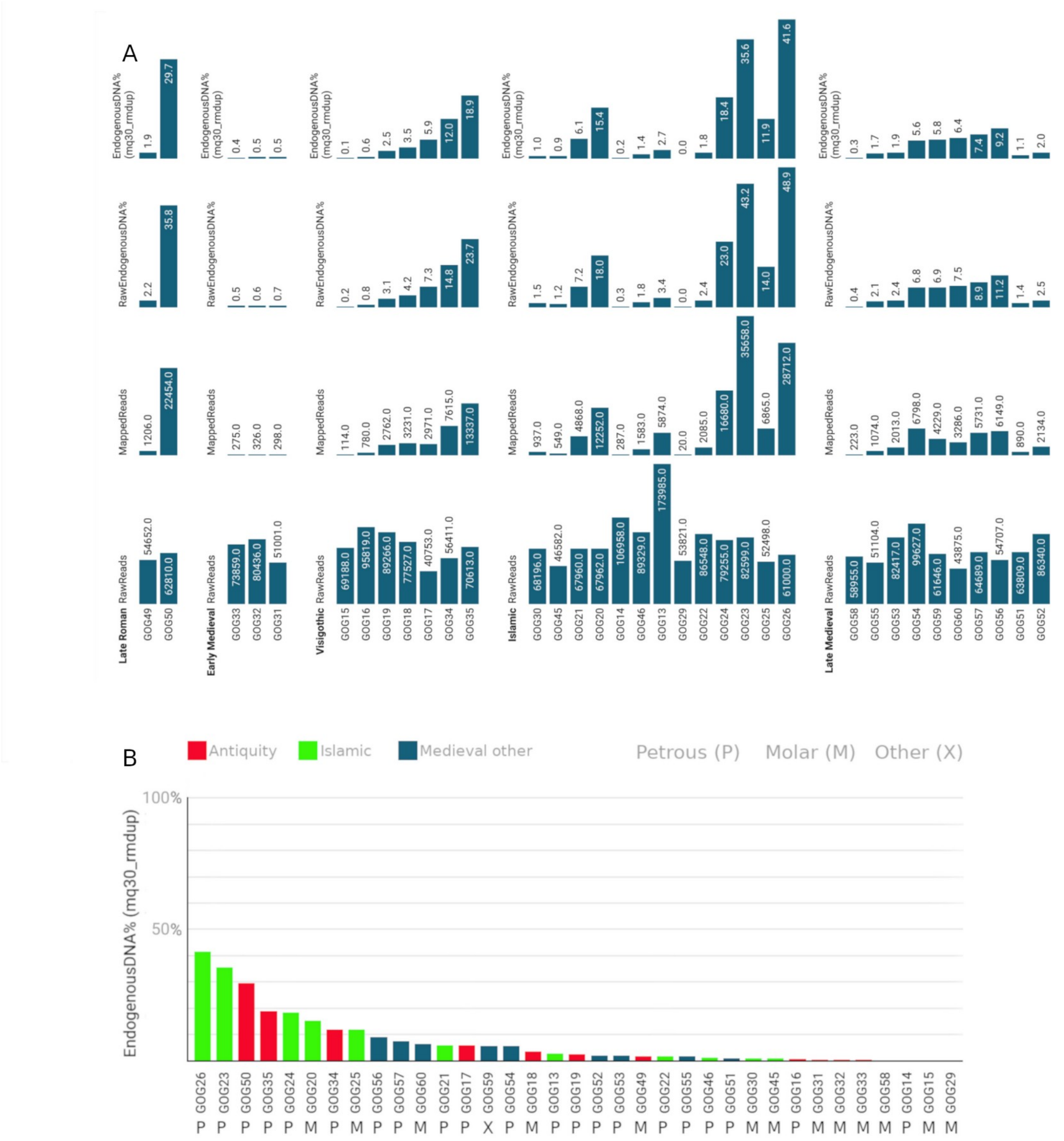


Figure S1: A) Results for all screened samples by sample and period. B) Adjusted endogenous DNA content after quality filters by sample, period and type of bone.

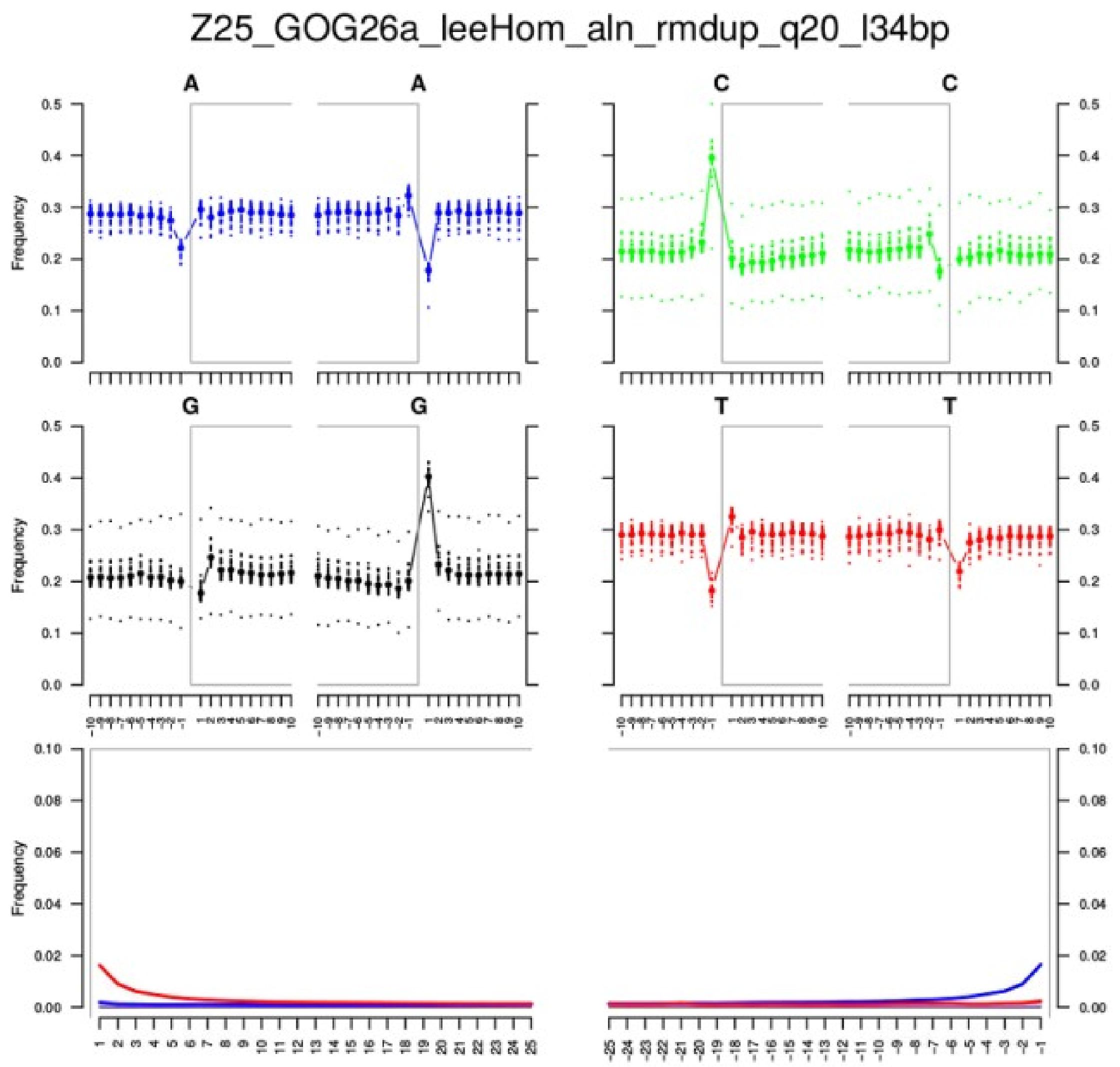


Figure S2: Example of reduced damage pattern in UDG-treated library. All libraries for all samples presented the same reduced pattern of damage typical of ancient DNA. 3bp were soft-clipped in all reads based on the damage patterns seen in the treated libraries. Full damage patterns had been previously confirmed in non-treated screening libraries.

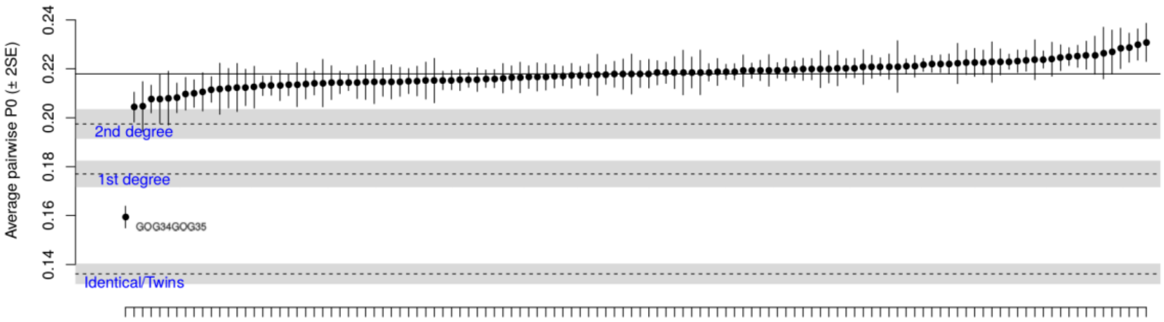


Figure S3: READ pairwise kinship coefficients combinations for all newly reported ancient samples.

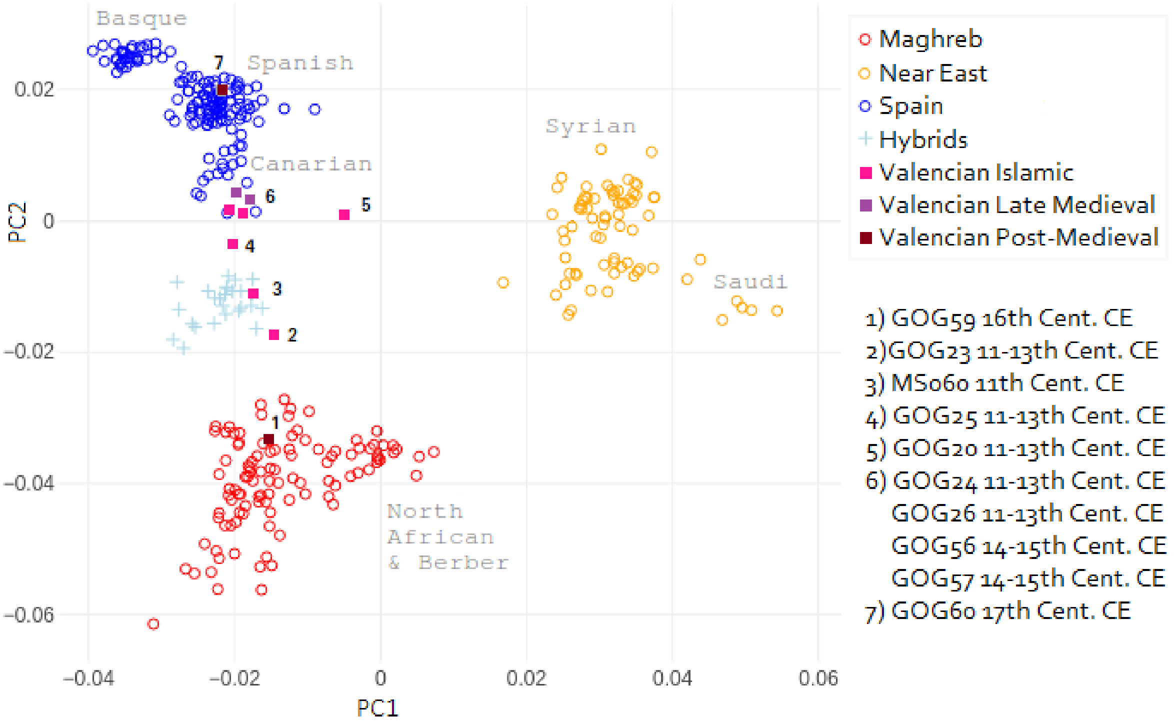

Figure S4: PCA zoom with modern populations merged from the HO and Arauna et al., (2017) datasets. Ancient medieval and simulated Spanish-Moroccan hybrid genomes projected.

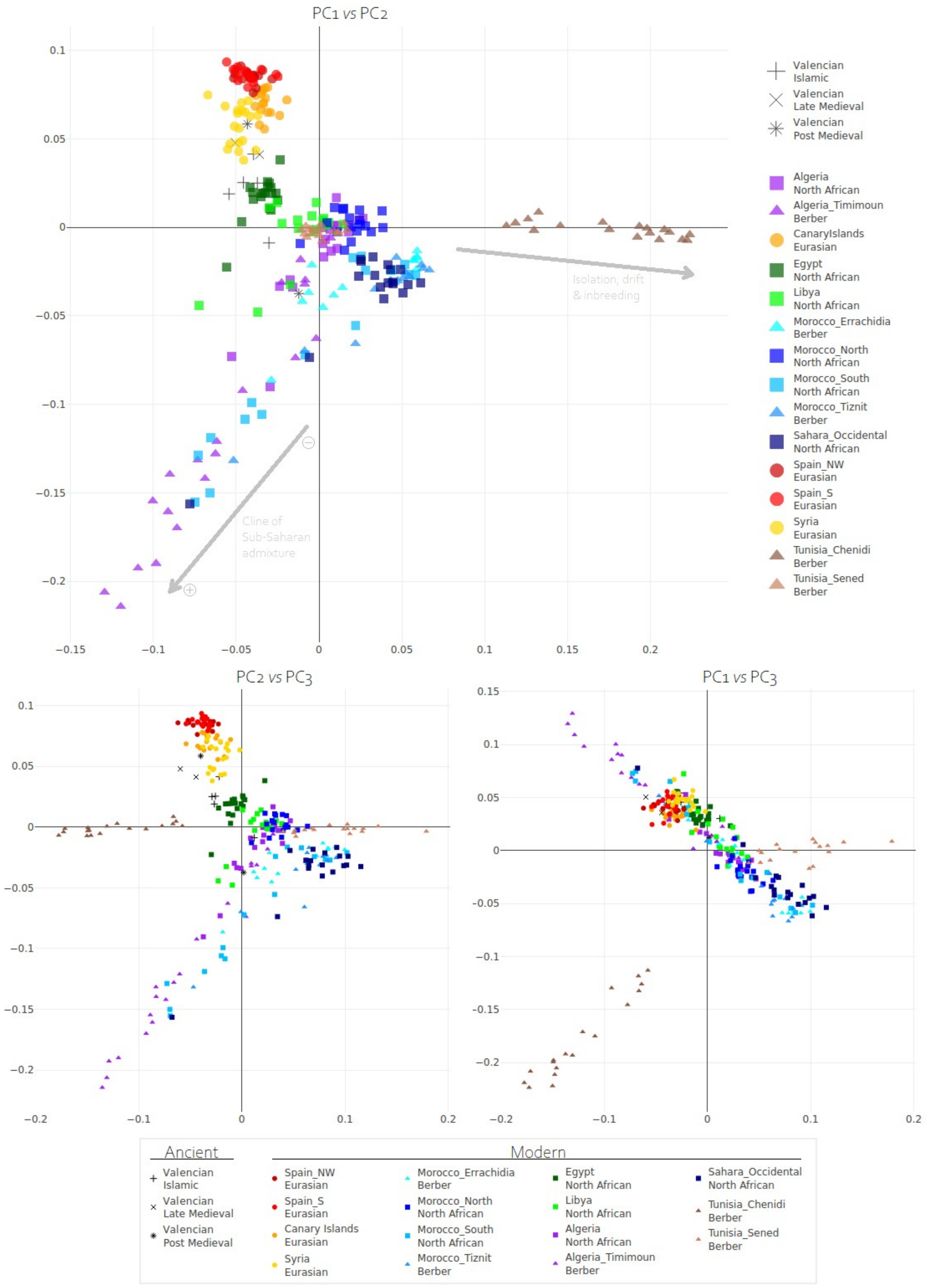

Figure S5: PCA with Arauna et al., (2017) dataset with the medieval samples projected.

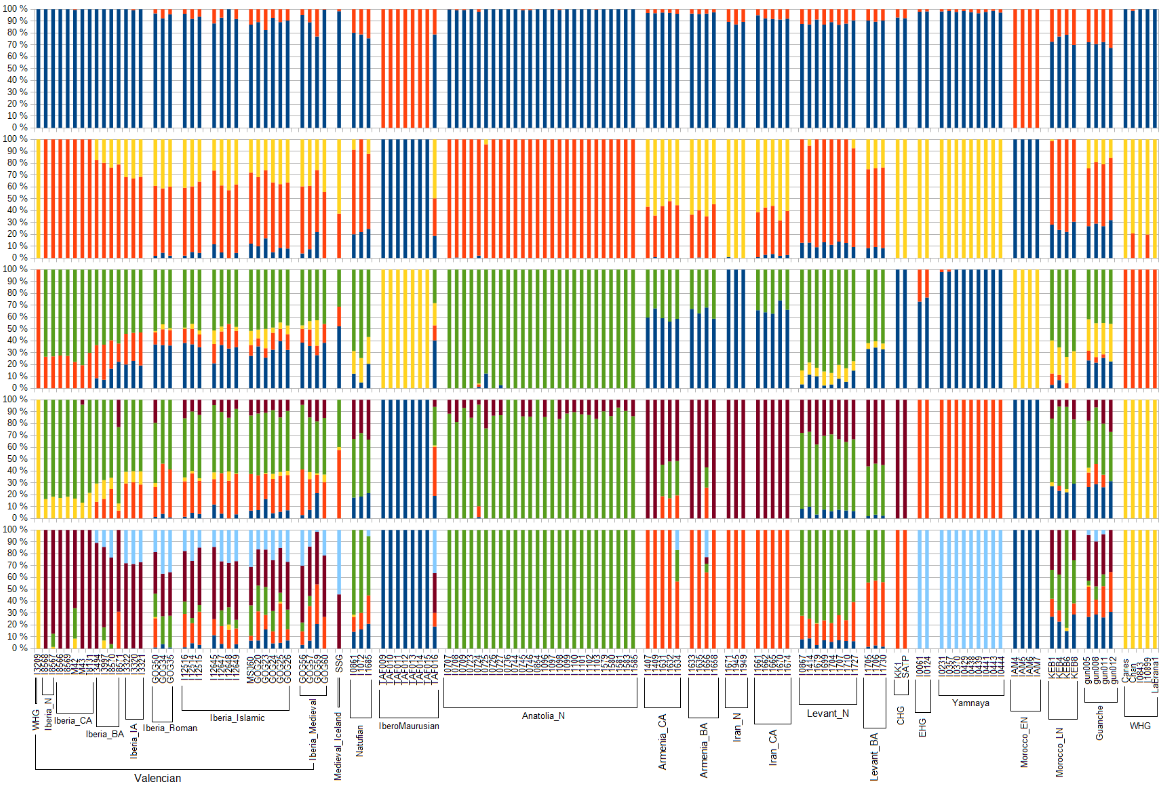


Figure S6: Unsupervised ADMIXTURE using only ancient populations projected in Figure 2A.

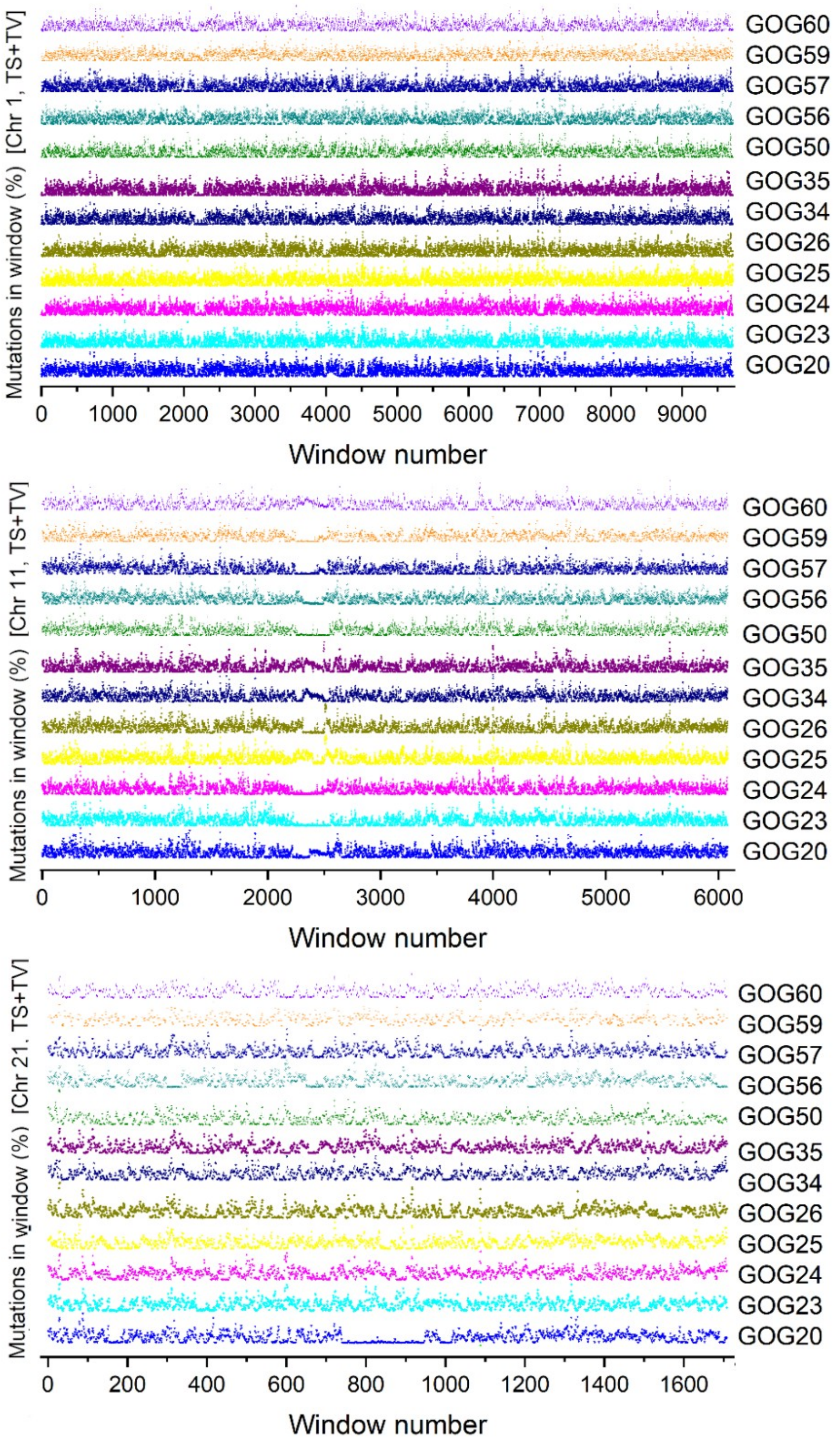
Mutations in window (%) [Chr 1, TS+TV]
GOG60
GOG59
GOG57
GOG56
GOG50
GOG35
GOG34
GOG26
GOG25
GOG24
GOG23
GOG20
0
1000
2000
3000
4000
5000
6000
7000
8000
9000
Window number
Mutations in window (%) [Chr 11, TS+TV]
GOG60
GOG59
GOG57
GOG56
GOG50
GOG35
GOG34
GOG26
GOG25
GOG24
GOG23
GOG20
0
1000
2000
3000
4000
5000
6000
Window number
Mutations in window (%) [Chr 21. TS+TV]
GOG60
GOG59
GOG57
GOG56
GOG50
GOG35
GOG34
GOG26
GOG25
GOG24
GOG23
GOG20
0
200
400
600
800
1000
1200
1400
1600
Window number

Figure S7: Sliding window of heterozygosity along the length of all imputed chromosomes (chr1, chr11, chr21) in the ancient genomes reported in this study.

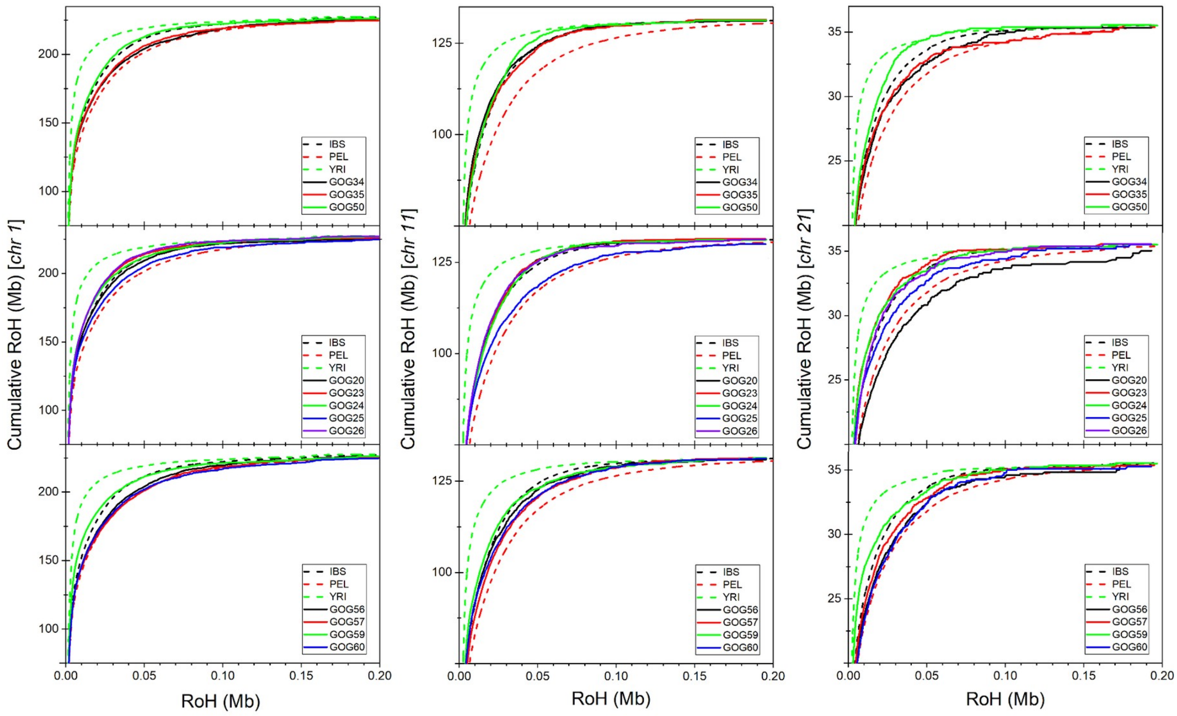


Figure S8: Cumulative organic ROH for all imputed chromosomes (chr1, chr11, chr21) in the ancient genomes reported in this study.

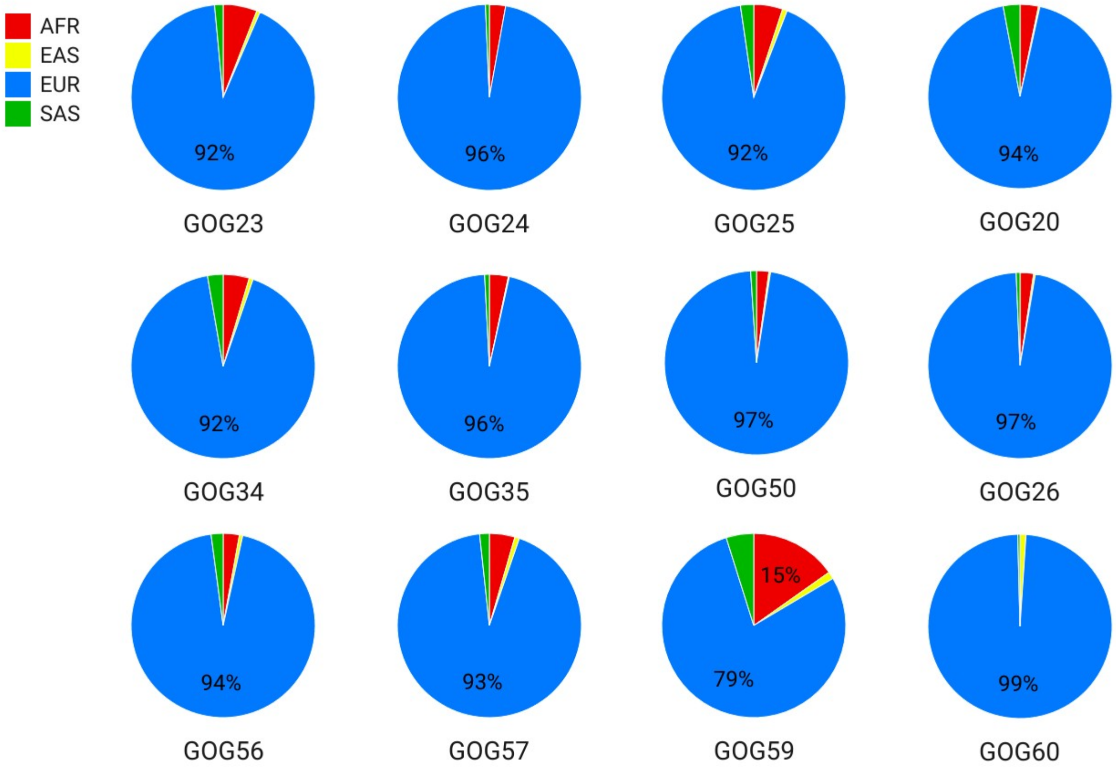


Figure S9: Ancestry inferred using RFMix combining all imputed chromosomes (chr1, chr11, chr21) in the ancient genomes reported in this study.

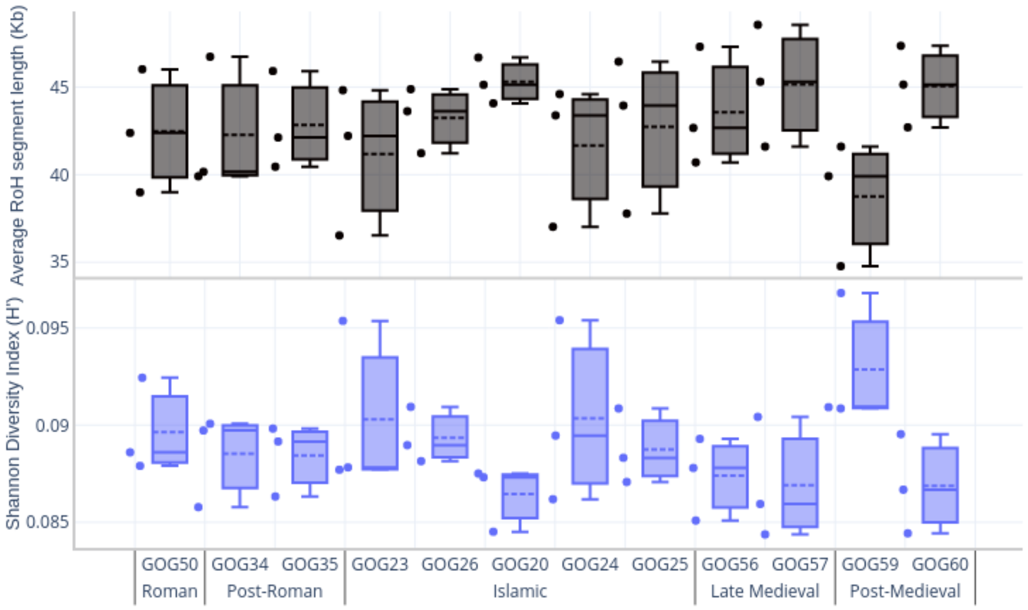


Figure S10: Average ROH length (Kb) in imputed chromosomes (top). Shannon diversity indexes calculated for each imputed chromosome (chr1, chr11, chr21) in the ancient genomes reported in this study using a number of variable positions.

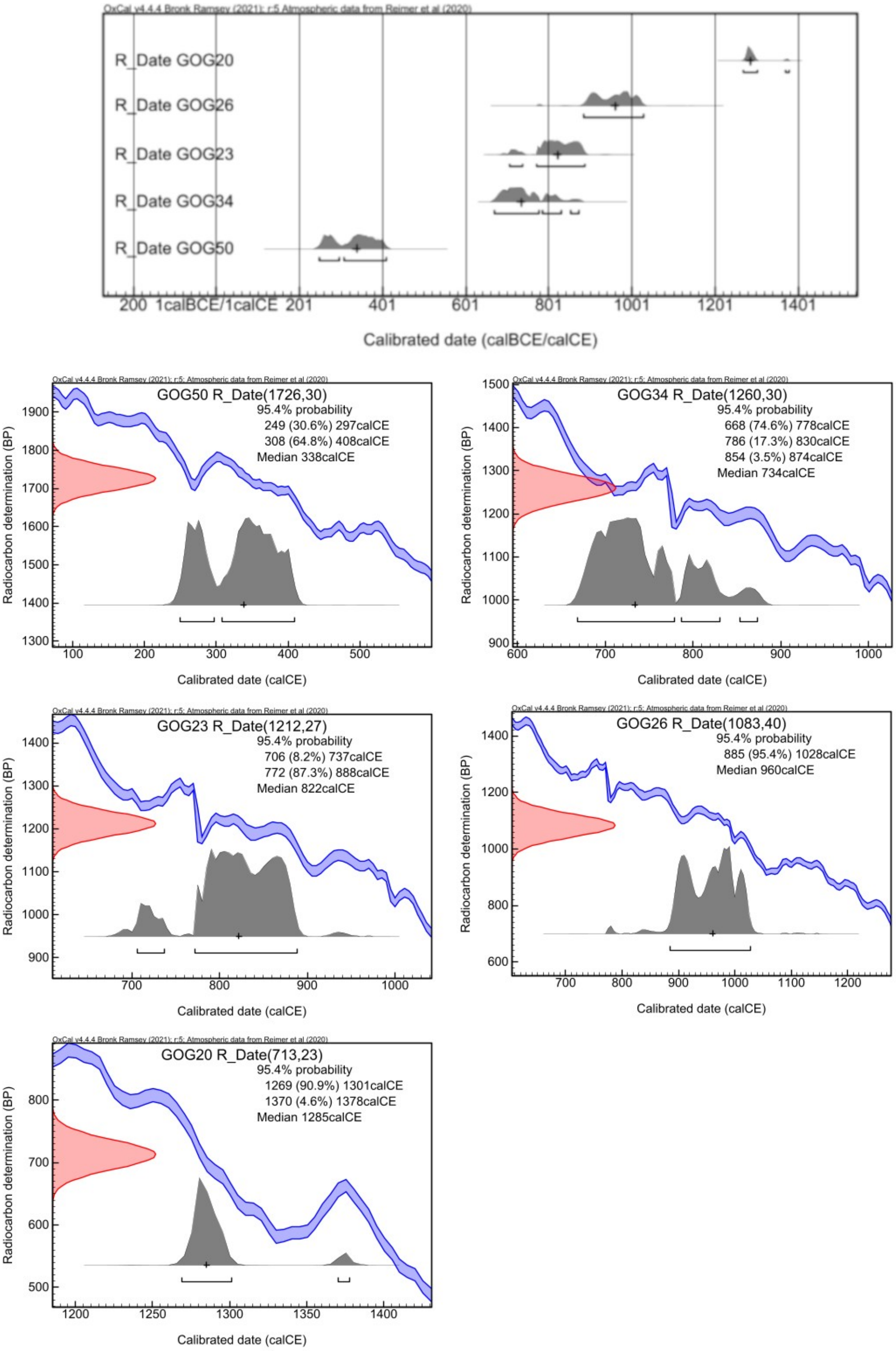


Figure S11: Radiocarbon dates obtained for a subset of five samples reported in this study.

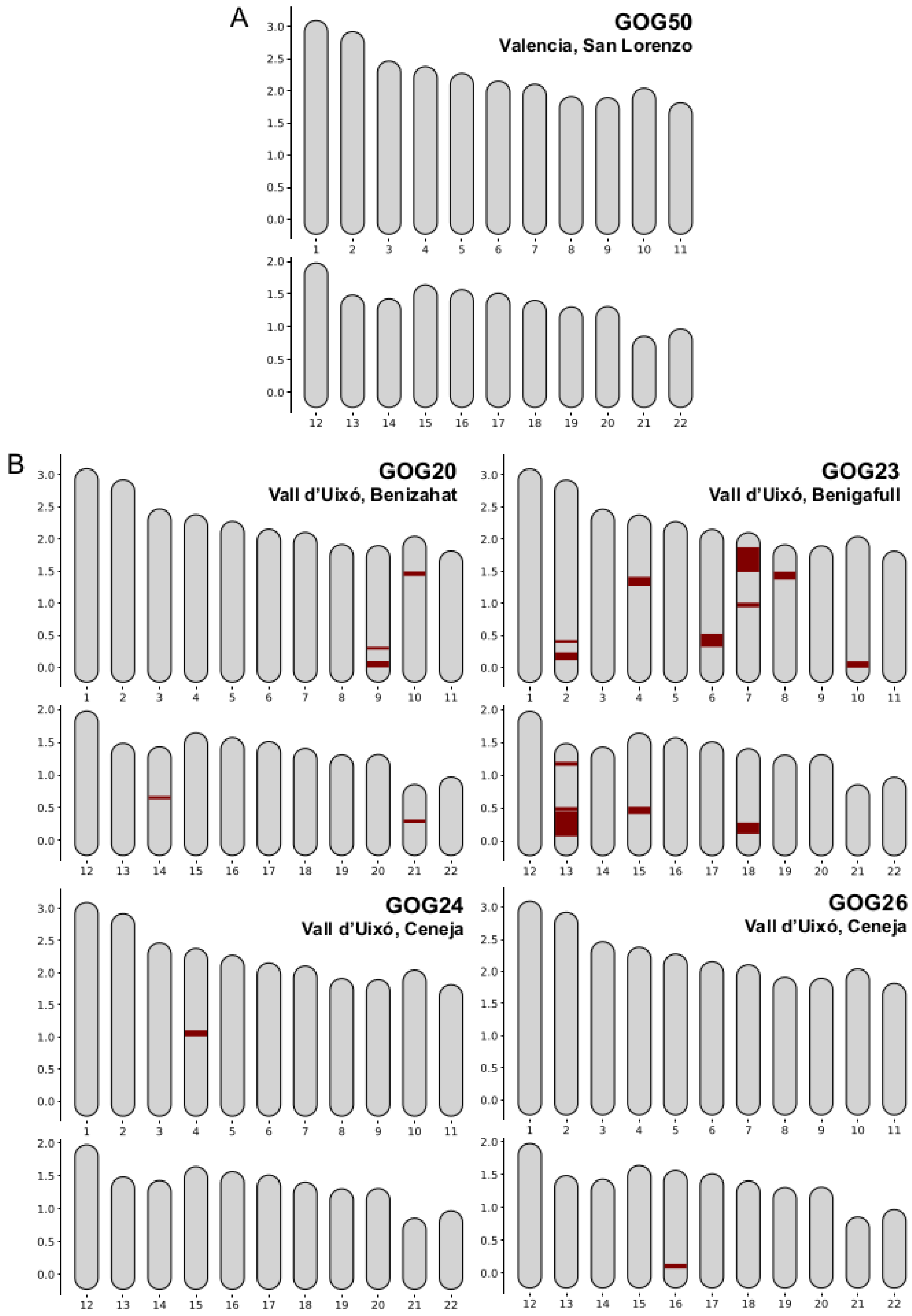


Figure S12: karyotypes with ROH by chromosome identified with hapROH.

### Supplementary References


1. Oteo-García G. Archaeogenetics of Southwest Europe [PhD Thesis]. University of Huddersfield; 2020. Available from: https://eprints.hud.ac.uk/id/eprint/35459
2. Olivé-Busom J, López-Costas O, Márquez-Grant N, Kirchner H. Estudio antropológico de las alquerías de Benizahat y Zeneta (Vall d’Uixó, Castellón). Una ventana a la vida rural andalusí. SAGVNTVM PLAV. 2021;53:193–212.
3. Oteo-García G, Alapont-Martín L, Pascual Beneyto J, Foody MGB, Yau B, Pala M, et al. Late Roman tombs at Sanxo Llop (Gandía, Valencia): Exogamy and kinship in a particular funerary structure. In: Death and the Societies of Late Antiquity: New methods, new questions? (Archéologies méditerranéennes). Aix-en-Provence: Presses universitaires de Provence; 2023. p. 119–27.
4. Silva M, Oteo-García G, Martiniano R, Guimarães J, von Tersch M, Madour A, et al. Biomolecular insights into North African-related ancestry, mobility and diet in eleventh-century Al-Andalus. Sci Rep. 2021;11(1):18121.
5. Pinhasi R, Fernandes D, Sirak K, Novak M, Connell S, Alpaslan-Roodenberg S, et al. Optimal ancient DNA yields from the inner ear part of the human petrous bone. PLoS ONE. 2015;10(6):e0129102.
6. Bronk Ramsey C. Bayesian analysis of radiocarbon dates. Radiocarbon. 2009;51(1):337–60.
7. Reimer P, Austin W, Bard E, Bayliss A, Blackwell P, Bronk Ramsey C, et al. The IntCal20 Northern Hemisphere Radiocarbon Age Calibration Curve (0–55 cal kBP). Radiocarbon. 2020;62(4):725–57.
8. Rohland N, Hofreiter M. Ancient dna extraction from bones and teeth. Nat Protoc. 2007;2(7):1756–62.
9. Yang DY, Eng B, Waye JS, Dudar JC, Saunders SR. Improved DNA extraction

from ancient bones using silica based spin columns. Am J Phys Anthropol. 1998;105(4):539–43.

10. MacHugh D, Edwards C, Bailey J, Bancroft D, Bradley D. The extraction and analysis of ancient DNA from bone and teeth: a survey of current methodologies. Anc Biomol. 2000;3.

11. Meyer M, Kircher M, Gansauge MT, Li H, Racimo F, Mallick S, et al. A high-coverage genome sequence from an archaic Denisovan individual. Science. 2012;338(6104):222–6.

12. Gamba C, Hanghøj K, Gaunitz C, Alfarhan AH, Alquraishi SA, Al-Rasheid KAS, et al. Comparing the Performance of Three Ancient DNA Extraction Methods for High-Throughput Sequencing. Mol Ecol Resour. 2016;16(2):459–69.

13. Cassidy LM, Martiniano R, Murphy EM, Teasdale MD, Mallory J, Hartwell B, et al. Neolithic and Bronze Age migration to Ireland and establishment of the insular Atlantic genome. Proc Natl Acad Sci. 2016;113(2):368–73.

14. Briggs AW, Stenzel U, Johnson PLF, Green RE, Kelso J, Prüfer K, et al. Patterns of damage in genomic DNA sequences from a Neandertal. Proc Natl Acad Sci U S A. 2007;104(37):14616–21.

15. Briggs AW, Stenzel U, Meyer M, Krause J, Kircher M, Pääbo S. Removal of deaminated cytosines and detection of in vivo methylation in ancient DNA. Nucleic Acids Res. 2009;38(6):e87.

16. Lindahl T. Instability and decay of the primary structure of DNA. Nature. 1993;362(6422):709–15.

17. Lindahl T. The Croonian Lecture, 1996: Endogenous damage to DNA. Philos Trans R Soc B Biol Sci. 1996;351(1347):1529–38.

18. Renaud G, Stenzel U, Kelso J. LeeHom: Adaptor trimming and merging for Illumina sequencing reads. Nucleic Acids Res. 2014;

19. García-Alcalde F, Okonechnikov K, Carbonell J, Cruz LM, Götz S, Tarazona S, et al. Qualimap: evaluating next-generation sequencing alignment data. Bioinformatics. 2012;28(20):2678–9.

20. Martiniano R, Cassidy LM, Ó'Maoldúin R, McLaughlin R, Silva NM, Manco L, et al. The population genomics of archaeological transition in west Iberia: Investigation of ancient substructure using imputation and haplotype-based methods. PLoS Genet. 2017;13(7).

21. Rohland N, Harney E, Mallick S, Nordenfelt S, Reich D. Partial uracil – DNA – glycosylase treatment for screening of ancient DNA. Philos Trans R Soc B Biol Sci. 2015;370(1660):20130624.

22. Jónsson H, Ginolhac A, Schubert M, Johnson PLF, Orlando L. MapDamage2.0: Fast approximate Bayesian estimates of ancient DNA damage parameters. In: Bioinformatics. 2013. p. 1682–4.

23. Malaspinas AS, Tange O, Moreno-Mayar JV, Rasmussen M, DeGiorgio M, Wang Y, et al. bammds: a tool for assessing the ancestry of low-depth whole-genome data using multidimensional scaling (MDS). Bioinforma Oxf Engl. 2014;30(20):2962–4.

24. Skoglund P, Storå J, Götherström A, Jakobsson M. Accurate sex identification of ancient human remains using DNA shotgun sequencing. J Archaeol Sci. 2013;40(12):1427–32.

25. McKenna A, Hanna M, Banks E, Sivachenko A, Cibulskis K, Kernytsky A, et al. The genome analysis toolkit: A MapReduce framework for analyzing next-generation DNA sequencing data. Genome Res. 2010;20(9):1297–303.

26. Thorvaldsdóttir H, Robinson JT, Mesirov JP. Integrative Genomics Viewer (IGV): High-performance genomics data visualization and exploration. Brief Bioinform. 2013;14(2):178–92.

27. Weissensteiner H, Pacher D, Kloss-Brandstätter A, Forer L, Specht G,

Bandelt HJ, et al. HaploGrep 2: mitochondrial haplogroup classification in the era of high-throughput sequencing. Nucleic Acids Res. 2016;44(W1):W58–63.

28. van Oven M. PhyloTree Build 17: Growing the human mitochondrial DNA tree. Forensic Sci Int Genet Suppl Ser. 2015;5:E392–4.

29. Ralf A, Montiel González D, Zhong K, Kayser M. Yleaf: Software for Human Y-Chromosomal Haplogroup Inference from Next-Generation Sequencing Data. Mol Biol Evol. 2018;35(5):1291–4.

30. Martiniano R, De Sanctis B, Hallast P, Durbin R. Placing Ancient DNA Sequences into Reference Phylogenies. Mol Biol Evol. 2022;39(2):msac017.

31. Kuhn JMM, Jakobsson M, Günther T. Estimating genetic kin relationships in prehistoric populations. PLoS ONE. 2018;13(4):e0195491.

32. Mallick S, Reich D. The Allen Ancient DNA Resource (AADR): A curated compendium of ancient human genomes. 2023.

33. Arauna LR, Mendoza-Revilla J, Mas-Sandoval A, Izaabel H, Bekada A, Benhamamouch S, et al. Recent Historical Migrations Have Shaped the Gene Pool of Arabs and Berbers in North Africa. Mol Biol Evol. 2017;34(2):318–29.

34. Patterson N, Price AL, Reich D. Population structure and eigenanalysis. PLoS Genet. 2006 Dec;2(12):2074–93.

35. Lazaridis I, Patterson N, Mittnik A, Renaud G, Mallick S, Kirsanow K, et al. Ancient human genomes suggest three ancestral populations for present-day Europeans. Nature. 2014;513:409–13.

36. Alexander DH, Novembre J, Lange K. Fast model-based estimation of ancestry in unrelated individuals. Genome Res. 2009;19(9):1655–64.

37. Green RE, Krause J, Briggs AW, Maricic T, Stenzel U, Kircher M, et al. A draft sequence of the neandertal genome. Science. 2010;328(5979):710–22.

38. Reich D, Thangaraj K, Patterson N, Price AL, Singh L. Reconstructing

Indian population history. Nature. 2009;461(7263):489–94.

39. Peter BM. Admixture, population structure, and f-statistics. Genetics. 2016;202(4):1485–501.

40. Patterson N, Moorjani P, Luo Y, Mallick S, Rohland N, Zhan Y, et al. Ancient admixture in human history. Genetics. 2012;192:1065–93.

41. Patterson N, Isakov M, Booth T, others. Large-scale migration into Britain during the Middle to Late Bronze Age. Nature. 2022;601:588–94.

42. Maples BK, Gravel S, Kenny EE, Bustamante CD. RFMix: a discriminative modeling approach for rapid and robust local-ancestry inference. Am J Hum Genet. 2013;93(2):278–88.

43. Cassidy LM, Maoldúin RÓ, Kador T, Lynch A, Jones C, Woodman PC, et al. A dynastic elite in monumental Neolithic society. Nature. 2020;582:384–8.
